# Patient-Derived Melanoma Organoids Preserve Tumor-Immune Heterogeneity and Reveal Context-Dependent Responses to Immune Checkpoint Blockade

**DOI:** 10.64898/2026.09.23.753873

**Authors:** Mamatha Serasanmbati, Jay Chadokiya, Michitaka Nakano, Max Julve, Pauline Funchain, Saurabh Sharma, Emma Wagner, Prabhjeet Singh, Calvin J. Kuo, Allison Betof, Amanda Kirane

## Abstract

**Background:** Experimental models that retain endogenous tumor–immune complexity are needed to investigate heterogeneous responses to immune checkpoint inhibitors (ICIs) in melanoma. We established patient-derived melanoma organoids (PDMOs) to examine retention of parental tumor cellular components and investigate patient-specific and context-dependent responses to checkpoint blockade.

**Methods:** Fresh melanoma specimens (n=50), including primary tumors, neoadjuvant-treated tumors and tumor-infiltrating lymphocyte (TIL)-associated samples were processed for Matrigel-embedded culture of patient-derived melanoma organoids (PDMOs). Characterization and functional studies were performed in subsets of established cultures. Immunofluorescence assessed tumor, stromal, immune, and checkpoint-marker expression in PDMOs and matched parental tissues. NGFR expression was evaluated in relation to organoid establishment and growth. Responses to anti-PD-1 alone or combined with anti-LAG-3 or anti-CTLA-4 therapies were evaluated using viability and morphological analyses, with selected models undergoing immune phenotyping and multiplex cytokine profiling. Available clinical outcomes were used for exploratory comparison. PDMOs and matched two-dimensional cultures were evaluated under 21% and 5% oxygen conditions.

**Results:** PDMOs were established from 30 of 50 specimens (60%) and retained melanoma, stromal, lymphoid, and myeloid components identified in matched parental tissues during early culture, together with immune checkpoint expression. Higher NGFR (CD271) expression was associated with greater organoid-forming capacity in an exploratory subset. Responses to checkpoint blockade varied across patient-derived models and regimens, with combination blockade not uniformly producing lower viability than PD-1 monotherapy. Ex vivo responses showed parallels with available clinical outcomes, with discordant cases also observed. In three selected PDMOs, divergent PD-1 responses were accompanied by differences in T-cell and myeloid representation, MHC-II and AXL expression, and inflammatory versus immunoregulatory cytokine profiles. Under normoxic conditions, PDMOs and matched two-dimensional cultures exhibited divergent treatment sensitivities, including clinically responding cases in which sensitivity was retained in PDMOs but attenuated in monolayers. Changing oxygen tension further altered checkpoint sensitivity within individual PDMOs, with the direction and magnitude of the observed shifts varying by model and regimen.

**Conclusions:** Early passage PDMOs retain key endogenous tumor, immune, and stromal components and support functional investigation of patient-derived melanoma biology. Their heterogenous responses across checkpoint regimens and oxygen conditions provide a platform for investigating how tumor-immune composition and environmental context influence treatment sensitivity.

## Introduction

Melanoma is the most aggressive form of skin cancer and accounts for the majority of skin cancer-related deaths worldwide. Its high degree of genetic and phenotypic heterogeneity, remarkable metastatic potential and capacity to evade immune surveillance contribute to poor clinical outcomes, particularly in patients with advanced disease. Although the introduction of immune checkpoint inhibitors (ICIs), including antibodies targeting programmed cell death protein-1 (PD-1), cytotoxic T-lymphocyte-associated protein-4 (CTLA-4), and more recently lymphocyte activation gene-3 (LAG-3), has revolutionized melanoma treatment and significantly improved long term survival, only a subset of patients achieves durable clinical responses. Approximately 30-40% of patients respond to PD-1 monotherapy, while many exhibit either primary or acquired resistance despite treatment with combination immunotherapy. Consequently, identifying patients most likely to benefit from ICIs remains a major challenge in precision oncology ^1–3^.

Current biomarkers used to predict immunotherapy response, including PD-L1 expression, tumor mutational burden and immune infiltration, have limited predictive accuracy when applied individually because melanoma progression is driven by complex interactions among tumor cells, stromal components and the immune microenvironment. Moreover, tumor heterogeneity and dynamic changes during therapy further complicate patient stratification. These limitations highlight the urgent need for physiologically relevant preclinical models capable of preserving patient-specific tumor architecture, cellular diversity, and immune composition to facilitate therapeutic prediction and mechanistic studies ^4,5^.

Conventional two-dimensional (2D) melanoma cell cultures have been helpful in identifying key oncogenic pathways but fail to reproduce the three-dimensional organization, extracellular matrix interactions and cellular heterogeneity present in native tumors. Similarly, patient-derived xenograft models, while retaining many tumor characteristics, are time-consuming, costly, and limited by the replacement of the human immune system with murine counterparts, restricting their application for immunotherapy studies. Consequently, these models often inadequately predict clinical responses to immune-based therapies ^6,7^.

Patient-derived organoids (PDOs) have emerged as promising three-dimensional ex vivo models that more faithfully recapitulate the histopathological architecture, genetic alterations, transcriptional programs, and phenotypic heterogeneity of their parental tumors. Generated directly from patient tumor tissue and cultured within extracellular matrix scaffolds, organoids preserve spatial organization, cell-cell interactions, and key microenvironmental features absent from conventional monolayer cultures. Importantly, patient-derived melanoma organoids (PDMOs) have demonstrated high molecular concordance with matched patient tumors and have shown considerable promise for high-throughput drug screening and personalized therapeutic testing ^8–10^. Recent studies further demonstrate that melanoma organoids can be co-cultured with autologous immune cells or stromal populations, providing physiologically relevant platforms for investigating tumor-immune interactions and evaluating immunotherapeutic strategies ^11–13^.

Melanoma is also characterized by pronounced phenotypic plasticity, whereby tumor cells dynamically transition between differentiated and dedifferentiated states associated with therapeutic resistance and disease progression. Regulators including microphthalmia-associated transcription factor (MITF), AXL, and nerve growth factor receptor (NGFR/CD271) have been implicated in melanoma stem-like properties, tumor initiation, and immune evasion. In parallel, the composition and functional state of infiltrating immune populations including cytotoxic T lymphocytes, macrophages, and antigen-presenting cells critically influence responses to checkpoint blockade. However, the extent to which patient-derived melanoma organoids preserve these cellular states and immune populations during long-term culture remains incompletely understood ^14–17^.

Despite growing interest in melanoma organoid technology, important knowledge gaps remain regarding their ability to faithfully retain the immune landscape of the parental tumor and to functionally predict patient-specific responses to immune checkpoint inhibition. Furthermore, relatively few studies have examined how tumor-intrinsic heterogeneity, three-dimensional architecture, and microenvironmental factors such as hypoxia collectively influence immunotherapy efficacy within organoid systems.

In this study, we established a biobank of patient-derived melanoma organoids generated from primary tumors, neoadjuvant-treated specimens, and tumors collected during tumor-infiltrating lymphocyte (TIL) therapy. We comprehensively characterized these organoids using histological, immunofluorescence, cytokine profiling, and immune phenotyping to determine their capacity to preserve tumor-immune heterogeneity. We further investigated the role of NGFR in organoid establishment and growth, evaluated functional responses to clinically relevant immune checkpoint inhibitor combinations, and examined the influence of three-dimensional architecture and oxygen tension on immunotherapy efficacy. Finally, we compared organoid responses with available clinical outcomes in an exploratory manner to estimate candidate viability threshold for prospective validation of the translational potential of patient-derived melanoma organoids as functional patient avatars for precision immuno-oncology.

## Materials and Methods

### Human melanoma specimen procurement

Fresh melanoma specimens were obtained from patients undergoing surgical resection or therapeutic procedures at Stanford University under an Institutional Review Board-approved tissue collection protocol (IRB #65607). Written informed consent was obtained from all participants before tissue collection. Samples included primary melanoma, metastatic melanoma, neoadjuvant pre-treatment tumors, post-treatment tumors, and tumor-infiltrating lymphocyte (TIL)-associated specimens. Clinical information including age, sex, disease stage, treatment history, and mutational status (including *BRAF* status) was recorded. All specimens were histopathologically confirmed as melanoma by a pathologist.

A total of 50 melanoma specimens were collected, of which 30 successfully generated patient-derived melanoma organoids (PDMOs). Clinicopathological characteristics of the study cohort are summarized in Table S1.

### Generation and culture of Patient Derived Melanoma Organoid (PDMOs)

Fresh tumor specimens were collected in RPMI-1640 medium (Gibco) supplemented with 10% fetal bovine serum (FBS; Gibco) and transported on ice for processing within 24. Organoids were established using previously described protocols with minor modifications ^19–20^. Briefly, tumor tissues were washed two to three times with Dulbecco’s phosphate-buffered saline (DPBS; Gibco) and surrounding necrotic, connective and vascular tissues were carefully removed. Samples were mechanically finely minced by using sterile scalpels and resulting cell suspension was centrifuged at 1,200 rpm for 5 minutes at 4°C, and pellet was resuspended in cold growth factor-reduced Matrigel (Corning, CLS356231) at a final concentration of 75%. Approximately 25-30 μL Matrigel droplets containing cells were plated in six-well culture plates and allowed to polymerize for 15-20 minutes at 37°C before adding of organoid culture medium.

The culture medium consisted of 40% Advanced DMEM/F12 (Gibco), 50% L-WRN conditioned medium (ATCC; containing Wnt3A, R-spondin-1, and Noggin), and 10% heat-inactivated FBS supplemented with 1× GlutaMAX, 1 mM HEPES, 10 mM nicotinamide, 1 mM N-acetylcysteine, 1× B27 supplement without vitamin A, 0.5 μM A83-01, 10 nM gastrin, 50 ng/mL epidermal growth factor (EGF), 50 ng/mL fibroblast growth factor-1 (FGF-1), and 1× penicillin-streptomycin. During the first two weeks of culture, Normocin (InvivoGen) was included to prevent microbial contamination. Organoids were maintained at 37°C in an incubator with 5% CO₂, and culture medium was replaced every three days. Organoids were typically established within 7-14 days and subsequently expanded for downstream analyses.

For passaging, organoids were incubated with TrypLE Express (Gibco) for 5-10 minutes at 37°C to generate small clusters, which were embedded in fresh Matrigel for continued expansion. Successfully established PDMOs demonstrated stable three-dimensional architecture and could be serially passaged. Long-term cryopreservation was performed using freezing medium containing 90% FBS and 10% dimethyl sulfoxide (DMSO), followed by storage in liquid nitrogen.

### Organoid morphology and growth analysis

Organoid morphology was monitored using brightfield microscopy throughout culture. Representative images were acquired on days 4, 7, 14, and 21 using an inverted microscope equipped with a Hamamatsu digital camera (Hamamatsu, Japan). Organoid diameter and growth kinetics were quantified using ImageJ software. Growth characteristics were additionally evaluated across serial passages (P1-P4) in representative organoids.

### Immunofluorescence staining

For immunofluorescence staining, PDMOs embedded in Matrigel were fixed in 4% paraformaldehyde for 45 minutes at room temperature and subsequently embedded in HistoGel™ (Thermo Fisher Scientific, HG-4000-012). Matched parental tissue and PDMOs were fixed in 70% ethanol for at least 2 hours, dehydrated through graded ethanol solutions, followed by isopropanol and acetone, and paraffin embedded. Sections were cut at 5 µm thickness. Immunofluorescence staining was performed using standard protocols. Sections were incubated overnight at 4°C with primary antibodies targeting α-SMA (Abcam, ab7817, 1:200), Melan-A (Abcam, ab210546, 1:300), CD45 (Cell Signaling Technology, 13917S, 1:250), CD3ε (Abcam, ab5690, 1:100), CD8 (Invitrogen, MA5-13473, 1:100), CD11C (Lunaphore, MR10070, 1:200), CD11B (Novus, NB110-89474, 1:250), CD271/NGFR (Invitrogen, 14-9400-82, 1:100), AXL (eBioscience, 14-940-73, 1:50), PD-1 (Lunaphore, MR10120, 1:100), PD-L1 (Abcam, ab228415, 1:200), CD4 (Lunaphore, MR10020, 1:100), MITF (Invitrogen, MA5-32554, 1:100), CD163 (Abcam, ab189915, 1:200), TFGβ (Invitrogen, MA5-16949, 1:100), CD68 (Abca, Ab955), LAG-3 (Cell Signaling Technology, 15372S, 1:50), MHC class II (Abcam, ab55152, 1:200). Following washing with PBS containing 0.1% Tween-20, sections were incubated with Alexa Fluor–conjugated secondary antibodies (anti-rabbit Alexa Fluor 647, anti-mouse Alexa Fluor 488; Invitrogen) for 1 hour at room temperature. Nuclei were counterstained with DAPI (1:1000), and images were acquired using a Zeiss LSM 710 confocal microscope. Marker expression was quantified as the percentage of positively stained cells across multiple representative regions of interest.

### Flow cytometry

Fresh tumor specimens were dissociated into single-cell suspensions using mechanical and enzymatic digestion. Cells were stained with fluorophore-conjugated antibodies against NGFR (CD271) according to the manufacturer’s protocol. Flow cytometric acquisition was performed on a BD FACSCanto II flow cytometer, and data were analyzed using FlowJo software. The frequency of NGFR-positive melanoma cells was quantified and correlated with organoid establishment efficiency and subsequent growth characteristics.

### Immune checkpoint inhibitor treatment

PDMOs were expanded for 7 days prior to ICI screening. Organoids were harvested by dissolving Matrigel using Cell Recovery Solution (Corning, CB40253) at 4 °C for 20-25 minutes with gentle pipetting. Suspensions were centrifuged at 250 × g for 5 minutes at 4 °C, washed and replated in 96-well plates at approximately 5,000 organoids per well in 5% Matrigel. Cultures were allowed to recover overnight before treatment.

Organoids were treated for seven days with nivolumab (anti-PD-1; 1 μM), relatlimab (anti-LAG-3; 1 μM), ipilimumab (anti-CTLA-4; 5 μM), or corresponding combination therapies (PD-1 + LAG-3 and PD-1 + CTLA-4). Untreated organoids served as controls. Cell viability was determined using the CellTiter-Glo® 3D Cell Viability Assay (Promega, G9681) according to the manufacturer’s instructions ^21–22^. Luminescence was measured using a BioTek microplate reader and normalized to untreated controls. Viability was normalized to untreated controls and expressed as percentage survival. All experiments were performed in biological triplicates, and data are presented as mean ± standard deviation.

### Hypoxia experiments

To investigate the influence of hypoxia on immunotherapy response, selected PDMOs and matched two-dimensional melanoma cultures were maintained under normoxia (21% O₂) or hypoxia (5% O₂) conditions throughout immune checkpoint inhibitor treatment. Cell viability following treatment was measured using the CellTiter-Glo® 3D assay and compared between oxygen conditions.

### Cytokine profiling

Conditioned media were collected from untreated organoids (day 0) and after seven days of immune checkpoint inhibitor treatment. Cytokine concentrations were quantified using a 48-plex human cytokine Luminex assay (Milliplex HCYTA-60K-PX48, EMD Millipore) at the Stanford Human Immune Monitoring Center.

Briefly, 25 μL of undiluted conditioned medium was incubated overnight with antibody-conjugated magnetic beads, followed by sequential incubation with biotinylated detection antibodies and streptavidin-phycoerythrin. Samples were analyzed using a Luminex FlexMap 3D instrument. All samples were analyzed in duplicate, and analytes with bead counts below 20 or below the detection limit across all samples were excluded from further analysis. The cytokine panel included inflammatory cytokines, chemokines, and angiogenic mediators including IFN-γ, TNF-α, IL-2, IL-6, IL-8, CXCL9, CXCL10, CCL2, CCL5, and VEGF-A, among others.

### Statistical analysis

Statistical analyses were performed using GraphPad Prism version 8. Data are presented as mean ± standard deviation (SD) unless otherwise indicated. Comparisons between two groups were performed using either two-tailed Student’s *t*-test or Mann–Whitney *U* test depending on data distribution. Comparisons among multiple groups were analyzed using one-way analysis of variance (ANOVA) followed by Tukey’s multiple-comparison test. Bonferroni correction was applied where appropriate for multiple testing, and categorical variables were compared using the χ² test. A two-sided *p* value <0.05 was considered statistically significant.

Organoid response was classified from CellTiter-Glo viability normalized to untreated control. A candidate viability threshold was estimated from the patient-regimen pairs with available clinical outcomes by identifying the range of thresholds that maximized discrimination between clinical responders and non-responders (57.3% to 72.3%); the midpoint (64.8%, rounded to 65%) was adopted as the candidate threshold. A model was classified as responder when the mean and all three replicate wells fell below 65% and the mean ± SD did not cross the threshold, as non-responder when the mean and all replicates fell above 65% under the same condition, and as indeterminate when the replicate range or mean ± SD spanned the threshold. The threshold is proposed for prospective validation and was not tested in an independent cohort. A two-sided p value <0.05 was considered statistically significant.

## Results

### Establishment of patient-derived melanoma organoids from freshly resected melanoma specimens

Fresh melanoma specimens were processed using a standardized workflow for patient-derived melanoma organoid (PDMO) generation along 2D cell culture (Figure 1A). Briefly, freshly resected tumors were mechanically dissociated, embedded in 75% Matrigel, and cultured under three-dimensional conditions to support organoid formation, expansion, cryopreservation, and downstream functional analyses including immunophenotyping, cytokine profiling, and immune checkpoint inhibitor (ICI) testing.

**Figure 1.**
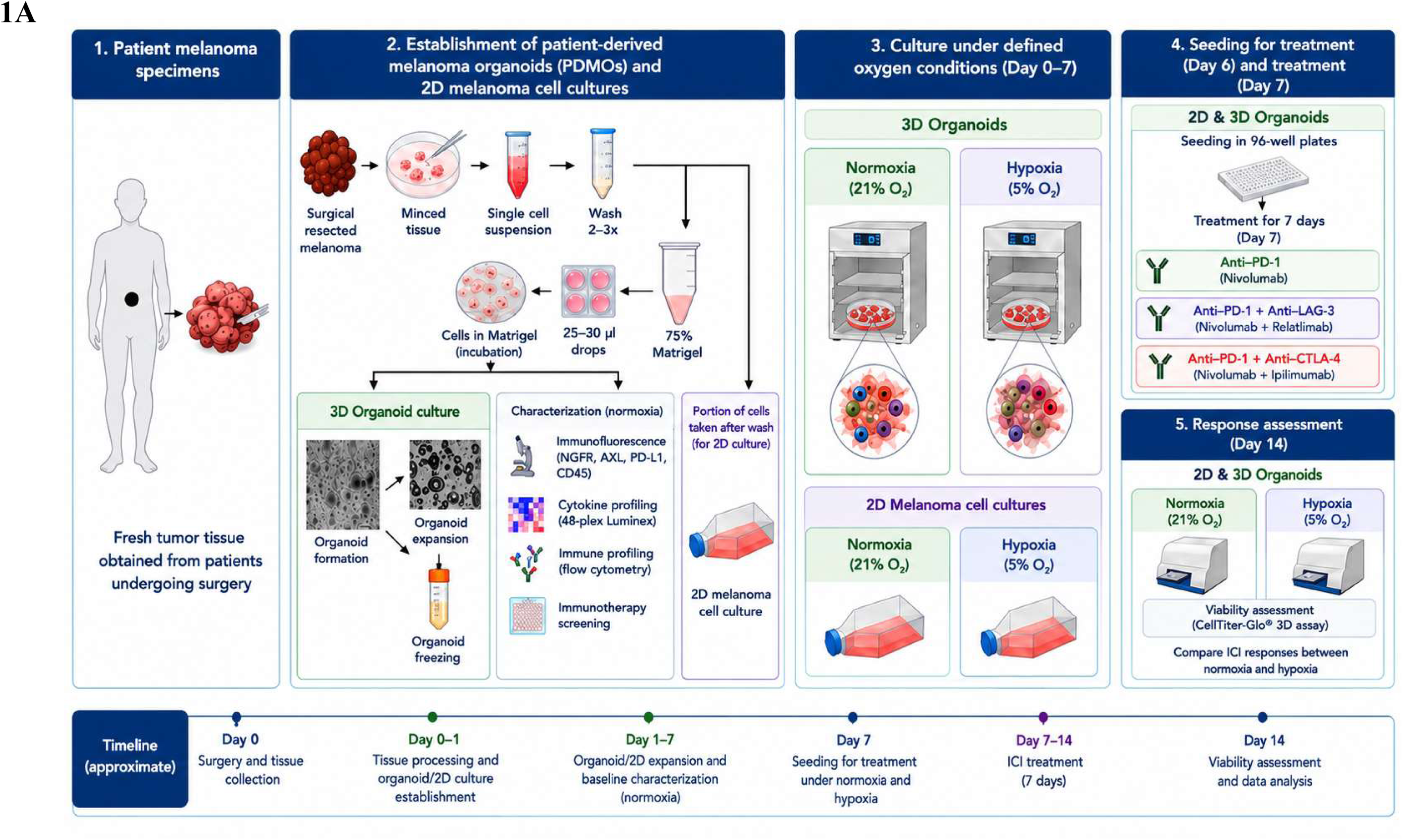
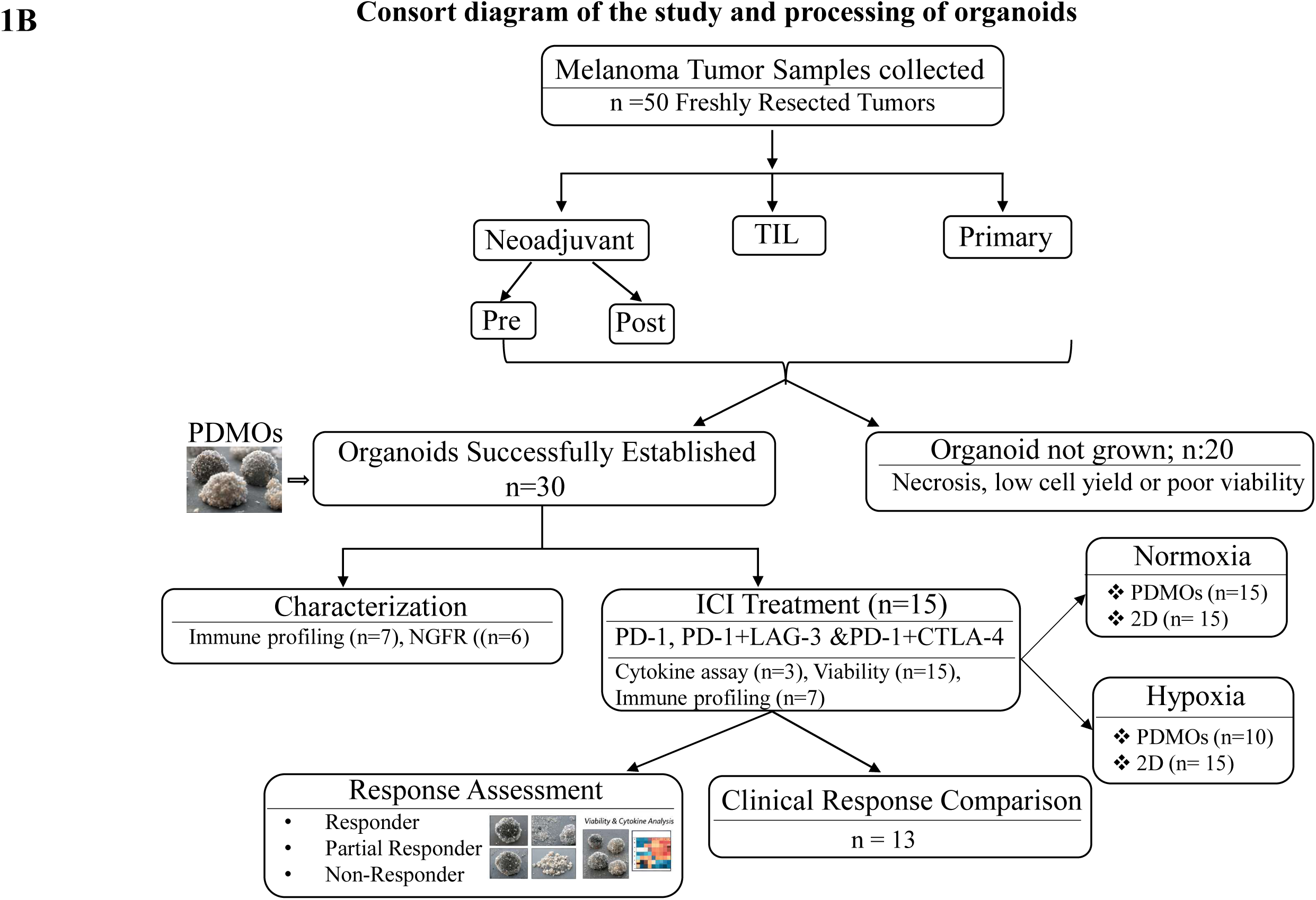
Experimental design for generation, characterization, clinical correlation and evaluating the impact of hypoxia on immune checkpoint inhibitor (ICI) efficacy in patient-derived melanoma organoids (PDMOs) and matched 2D melanoma cultures. **(A)** Fresh melanoma tissue obtained from patients undergoing surgical resection was mechanically dissociated to generate a single-cell suspension. Following washes, a portion of cells was collected for establishment of matched 2D melanoma cultures, while the remaining cells were embedded in 75% Matrigel as 25–30 μL drops for generation of patient-derived melanoma organoids (PDMOs). Organoids were expanded and characterized . Matched 3D organoids and 2D melanoma cultures were subsequently maintained under normoxic (21% O₂) or hypoxic (5% O₂) conditions. On day 7, cultures were seeded for treatment and exposed for 7 days to anti-PD-1 (nivolumab), anti-PD-1 + anti-LAG-3 (nivolumab + relatlimab), or anti-PD-1 + anti-CTLA-4 (nivolumab + ipilimumab). On day 14, treatment response was assessed by CellTiter-Glo® 3D viability assay under both oxygen conditions. The experimental timeline is shown from tissue collection and processing (day 0), organoid/2D culture establishment and baseline characterization (days 0-7), treatment seeding (day 7), ICI treatment (days 7-14), and final viability assessment and data analysis (day 14). **(B)** In parallel, a total of 50 freshly resected melanoma samples, including neoadjuvant (pre and post treatment), tumor-infiltrating lymphocyte (TIL), and primary tumors, were processed for organoid generation. Organoids were successfully established from 30 samples, while 20 failed due to necrosis, low cell yield, or poor viability. Established PDMOs underwent immune profiling, NGFR expression analysis and cytokine assays, followed by treatment with immune checkpoint inhibitors (PD-1, PD-1 + LAG-3 and PD-1 + CTLA-4). Treatment responses were assessed using viability and cytokine analyses and classified as responder, partial responder, or non-responder. **Abbreviations:** 2D:two-dimensional; 3D:three-dimensional; CTG: CellTiter-Glo; CTLA-4:cytotoxic T-lymphocyte-associated protein 4; ICI: immune checkpoint inhibitor; IF: immunofluorescence; LAG-3: lymphocyte activation gene-3; NGFR: nerve growth factor receptor; O₂: oxygen; PD-1: programmed cell death protein 1; PD-L1: programmed death-ligand 1; PDMO: patient-derived melanoma organoid.

A total of 50 freshly resected melanoma specimens representing primary tumors, neoadjuvant pre- and post-treatment tumors, and tumor-infiltrating lymphocyte (TIL)-associated samples were processed for organoid generation (Figure 1B). Stable organoids were successfully established from 30 specimens, corresponding to an overall establishment rate of 60%, whereas organoid generation failed in 20 specimens because of extensive necrosis, low cellularity, or poor tissue viability or loss of stemness.

Following embedding in Matrigel, cellular aggregates became apparent within 5-7 days and progressively developed into three-dimensional organoids over 7-14 days. Established PDMOs exhibited heterogeneous morphologies, including compact spheroidal, multilobulated, and irregular structures, reflecting the histological diversity of the originating tumors. Organoids could be expanded by serial passaging and successfully cryopreserved without obvious loss of viability or structural integrity after recovery, enabling the establishment of a living biobank for downstream studies.

Successfully established PDMOs were subsequently subjected to molecular and immune characterization, including immunofluorescence analysis, NGFR expression profiling, multiplex cytokine analysis, and functional evaluation of responses to PD-1, PD-1/LAG-3, and PD-1/CTLA-4 blockade (Figure 1B). Collectively, these findings demonstrate the feasibility of establishing patient-derived melanoma organoids from fresh melanoma specimens and support their application as an ex vivo platform for immune profiling and personalized immunotherapy testing.

### Morphological and growth dynamics of PDMOs

Brightfield imaging demonstrated the progressive formation and maturation of PDMOs over a 21-day culture period (Figure 2A). Small, loosely aggregated cell clusters were observed by Day 4, which increased in size, cellular density and structural organization by Day 7. By Day 14, mature organoids with compact cellular architecture had formed, while by Day 21, large and complex organoids exhibiting heterogeneous internal morphology were evident, consistent with continued proliferation and tissue maturation. Distinct morphological differences were observed among patient derived cultures, reflecting inter-patient heterogeneity in organoid forming capacity and tumor growth characteristics. Notably, organoids derived from Pt-37 and Pt-54 displayed robust growth, characterized by progressive increases in diameter, spherical morphology and greater structural complexity, whereas Pt-10 organoids remained smaller, less organized and exhibited limited expansion throughout the culture period.

**Figure 2:**
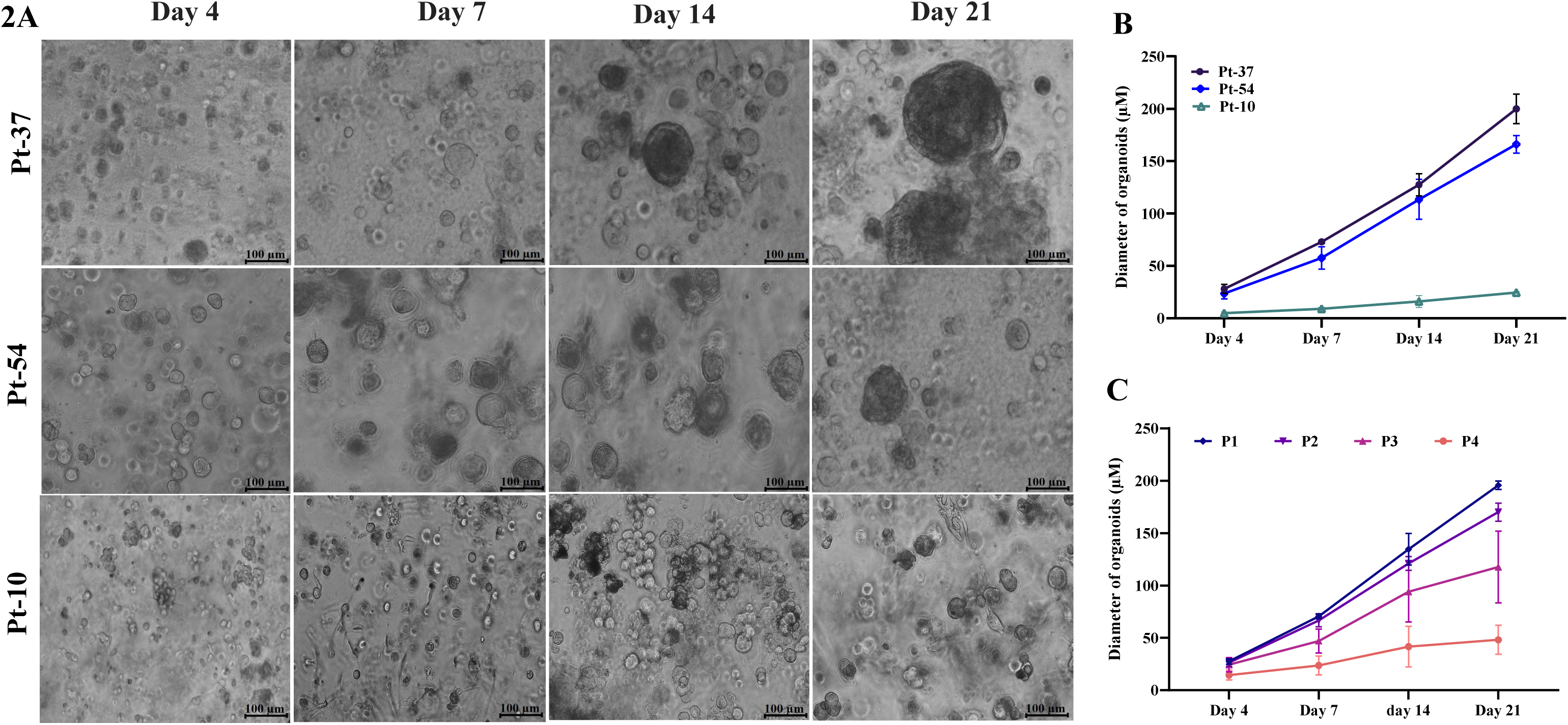
Morphological and growth dynamics of PDMOs. **(A)** Representative brightfield images of PDMOs derived from three melanoma patients (Pt-37, Pt-54, and Pt-10) cultured for 4, 7, 14, and 21 days. Pt-37 and Pt-54 progressively increased in size and structural complexity, developing well-defined spherical morphologies by Day 21. In contrast, Pt-10 formed smaller and less organized structures with minimal growth over time. Scale bars, 100 µm. **(B)** Quantitative analysis of organoid diameter for all PDMOs at different time points. At 21-day culture period showing patient-specific growth kinetics. Pt-37 and Pt-54 exhibited significantly greater expansion than Pt-10, reflecting inter-patient variability in organoid-forming capacity. Data are presented as mean ± SD. **(C)** Pt-37 organoid growth kinetics were observed in four sequential passages (P1-P4). Early-passage organoids (P1-P2) demonstrated higher proliferative capacity, whereas later passages (P3-P4) showed reduced growth rates, indicating gradual decline in expansion potential during prolonged culture, P4 showing the lowest increase in organoid diameter by Day 21. Data represent mean ± SD, n = 3 replicates per time point.

**Figure 2.**
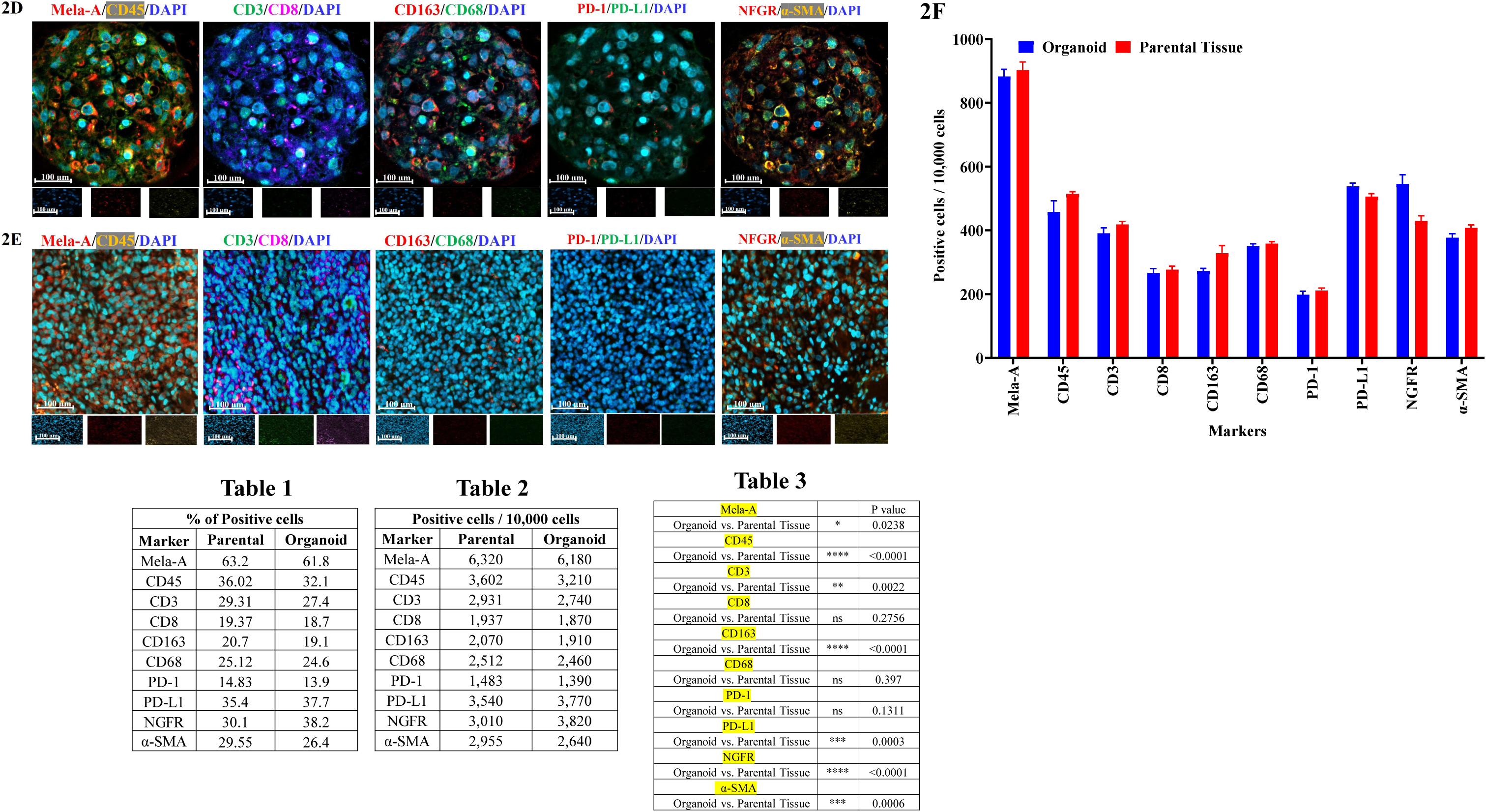
Immunofluorescence characterization proves preservation of tumor and immune heterogeneity in PDMOs. **(D)** Representative immunofluorescence images of PDMOs showing expression of melanoma lineage markers (Melan-A), phenotypic plasticity markers (AXL, NGFR), immune cell markers (CD45, CD3, CD8, CD68, CD163), stromal marker (α-SMA), and immune checkpoint molecules (PD-1 and PD-L1). Nuclei were counterstained with DAPI. Scale bars, 100 μm. **(E)** Corresponding immunofluorescence images of matched parental melanoma tissues demonstrating preservation of tumor, immune and immune checkpoint marker expression patterns in comparison with the derived organoids. **(F)** Quantitative comparison of marker-positive cells in matched parental tumors and PDMOs. Comparable expression profiles were observed across melanoma lineage, immune and immune checkpoint markers, with maintenance of Melan-A, CD45, CD68, CD163, α-SMA, PD-1, and PD-L1 expression. NGFR expression was modestly enriched in organoids, whereas slight reductions were observed in CD3⁺ and CD8⁺ T-cell populations. Data are presented as mean ± SD. Data represent n=7 (3 ROI quantification per organoid). Table 1-3: Quantification of % of positive cells on Immunofluorescence in Organoid vs matched Parental tissue and their p values.

Quantitative analysis confirmed a time-dependent increase in organoid diameter across all cultures, although growth kinetics varied considerably between patients (Figure 2B). Pt-37 and Pt-54 showed the greatest expansion, with mean organoid diameters increasing from approximately 40–50 μm on Day 4 to 180-220 μm by Day 21. In contrast, Pt-10 organoids exhibited only modest growth, reaching approximately 40-450 μm by Day 21, indicative of reduced organoid-forming efficiency and proliferative capacity.

To evaluate long-term culture stability, serial passaging experiments were performed using Pt-37 organoids (Passages 1-4) (Figure 2C). Early-passage organoids (P1-P2) retained strong proliferative capacity and demonstrated sustained increases in organoid diameter over time. In contrast, later passages (P3-P4) exhibited progressively reduced expansion kinetics, smaller organoid diameters, and diminished growth by Day 21, suggesting a gradual decline in proliferative potential during prolonged in vitro culture. Collectively, these findings demonstrate that PDMOs recapitulate patient-specific growth behavior while maintaining expansion capacity across multiple passages, although prolonged serial passaging is associated with reduced proliferative activity.

### PDMOs preserve tumor-intrinsic and immune heterogeneity of matched parental melanoma tissues

To determine whether PDMOs retained the cellular complexity of the original tumors, immunofluorescence was performed on matched parental melanoma tissues and organoids using markers of melanoma lineage, differentiation state, immune cell populations, stromal components, and immune checkpoint pathways (Figure 2D-F). Organoid establishment was successful in approximately 60% of patient samples, with reduced efficiency primarily observed in tumors exhibiting extensive therapy-associated necrosis. Established organoids remained viable for more than month but we noticed preserving key cellular and stromal components of the native tumor microenvironment until day 24 under our lab conditions.

PDMOs faithfully maintained melanoma lineage heterogeneity, demonstrating robust expression of the melanocytic marker Melan-A together with the lineage-specific transcription factor MITF and the dedifferentiation-associated markers AXL and NGFR (Figure 2D,E,S1A). The coexistence of MITF and AXL (Figure S1A) was high in tumor cell populations within individual organoids reflects the phenotypic plasticity characteristic of melanoma and closely mirrored the spatial distribution observed in matched parental tissues (Figure S1B). In addition to preserving tumor-intrinsic heterogeneity, PDMOs retained diverse immune cell populations, including CD45, CD3, CD4, CD8 T cells, CD68 macrophages, CD163 M2-like macrophages, and CD11B/CD11Cmyeloid cells. Stromal α-SMA-positive cells were also maintained, indicating preservation of essential components of the tumor microenvironment. Importantly, both PD-1 and PD-L1 (Figure 2D,E) remained detectable within organoid cultures, demonstrating conservation of clinically relevant immune checkpoint signaling pathways.

Quantitative comparison of matched parental tumors and organoids showed small absolute differences for most markers (Figure 2F, Table 1-3, S1C, Table S4A-C). Differences were within approximately 4% for Melan-A (61.8% in organoids vs. 63.2% in parental tissue), CD45 (32.1% vs. 36.02%), CD3, CD8, CD68, CD163, PD-1, PD-L1, α-SMA, AXL, CD4, TGFβ and CD11B, although several of these differences reached statistical significance within ROI-level replication. Larger differences were observed for MITF (22.9% vs. 28.4%) and CD11C (11.6% vs. 19.6%), which were lower in organoids, and for NGFR (38.2% vs 30.1%), which was higher, suggesting relative enrichment of dedifferentiated melanoma cells during ex vivo culture. Collectively, these findings demonstrate that PDMOs retain, during early culture, the principal tumor, immune, and stromal populations detected in matched parenteral tissue, with quantitative differences for a subset of markers.

### NGFR expression correlates with melanoma organoid-forming capacity

To study whether NGFR (CD271) expression is associated with melanoma organoid establishment, NGFR levels were assessed in six patient-derived melanoma specimens by immunofluorescence and flow cytometry (Figure 3A-C). NGFR expression exhibited marked inter-patient heterogeneity at both the tissue and single-cell levels. Immunofluorescence demonstrated abundant NGFR-positive cells in Pt-37, Pt-38 and Pt-164, whereas Pt-166, Pt-112, and Pt-65 showed minimal staining (Figure 3A). Flow cytometric analysis confirmed this variability, with NGFR-positive populations ranging from 0.38% to 89.1% across samples (Figure 3B). Notably, specimens containing higher proportions of NGFR-positive cells consistently generated robust, long-term organoid cultures, whereas tumors with low NGFR expression exhibited slow organoid growth or failed to establish organoids (Figure 3C). Specifically, Pt-37 (89.1%), Pt-38 (29.8%), and Pt-164 (11.9%) produced well-growing organoids, while Pt-112 (6.3%) generated slow-growing organoids and Pt-163 (1.16%) and Pt-166 (0.38%) failed to sustain organoid formation. In this exploratory series of six specimens, higher NGFR positivity accompanied more robust organoid growth; a larger series with prespecified growth criteria will be required to test NGFR as a predictor of organoid establishment.

**Figure 3.**
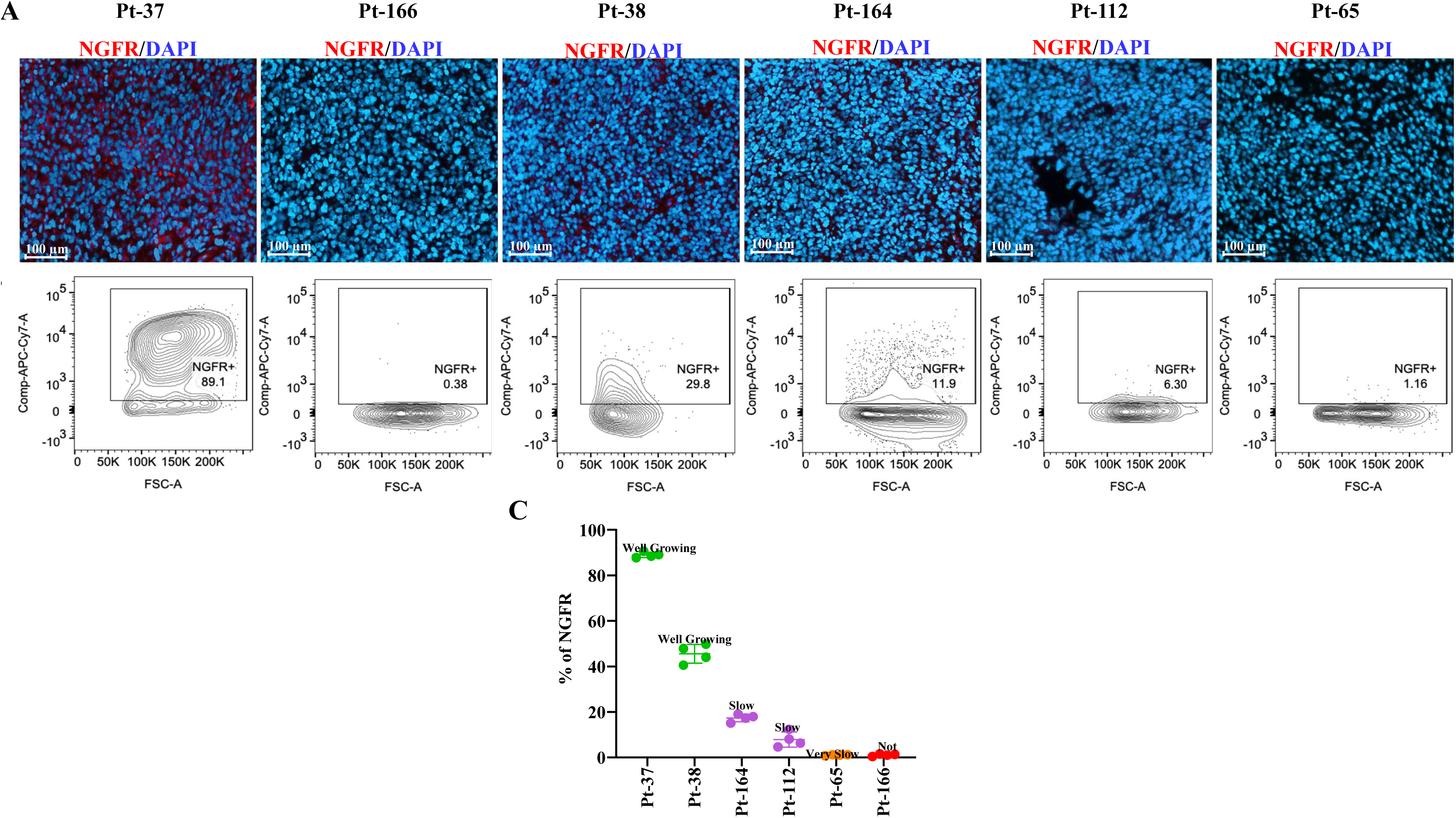
NGFR expression is associated with melanoma organoid-forming capacity: **(A)** Representative immunofluorescence images of melanoma specimens stained for NGFR (red) with DAPI nuclear counterstaining (blue). Scale bars, 100 μm. **(B)** Representative flow cytometry plots showing the frequency of NGFR-positive melanoma cells in each patient specimen. **(C)** Quantification of NGFR-positive cells in relation to organoid growth phenotype. Samples with higher NGFR expression generated robust organoids, whereas specimens with low NGFR positivity exhibited slow organoid growth or failed to establish organoid cultures, demonstrating a positive association between NGFR abundance and melanoma organoid-forming capacity.

### PDMOs recapitulate heterogeneous clinical responses to immune checkpoint inhibition and reveal distinct immune phenotypes associated with therapeutic sensitivity

To evaluate the utility of patient-derived melanoma organoids (PDMOs) as functional platforms for immunotherapy response prediction, organoids were treated with anti-PD-1 monotherapy or in combination with anti-LAG-3 or anti-CTLA-4 (Figure 4A-C). Functional viability assays demonstrated marked inter-patient heterogeneity, with organoids segregating into responder (R), Intermediate and non-responder (NR) phenotypes across treatment regimens. These results are consistent with clinical observations that a subset of melanoma tumors exhibit anti-PD-1 sensitivity due to pre-existing T-cell infiltration and preserved antigen presentation machinery ^23, 24^. The combination lowered viability relative to PD-1 alone in seven of fifteen models (Pt-53 most notably, 92.2% to 34.3%), was similar or higher in eight, and means were 61.3% vs 61.1%; anti-CTLA-4 lowered viability in four of fifteen (Supplemental Table 3A). The regimen associated with lowest viability therefore varied by model. Heatmap visualization further highlighted distinct patient-specific treatment signatures, demonstrating that individual organoids exhibited preferential sensitivity to different checkpoint inhibitor combinations. The primary objective of the clinical comparison was to estimate where a viability threshold should sit for prospective validation. Among 13 patients with available clinical outcomes (17 patient-regimen pairs), organoid calls at the candidate 65% threshold agreed with clinical response in 14 pairs, disagreed in 2 (Pt-54, PD-1+CTLA-4;) Pt-63, PD-1), and were intermediate in one (Pt-159, PD-1+CTLA-4) (Figure 4A-C & Table 4). Because the threshold was derived from these same outcomes, this agreement describes the fit of the candidate threshold rather than its predictive accuracy.

**Figure 4.**
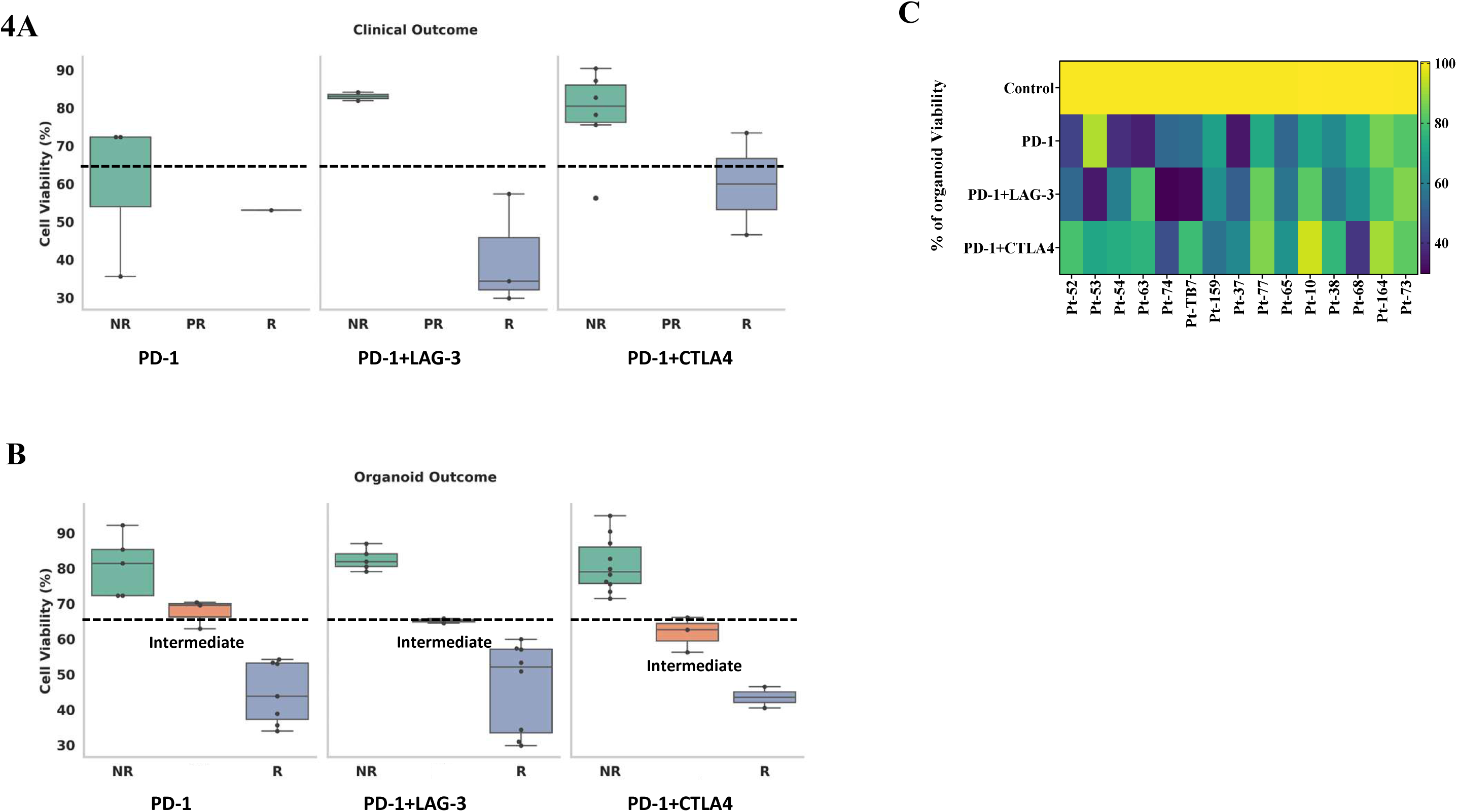
Patient-derived melanoma organoids recapitulate heterogeneous responses to immune checkpoint inhibition and reveal immune phenotypes associated with therapeutic sensitivity. **(A)** Clinical responder and non-responder specimen treatment pairs demonstrated significantly different ex vivo viability distributions. Exploratory threshold analysis identified equivalent maximal discrimination for thresholds between 57.3% and 72.3% viability; the midpoint of this interval (64.8%, rounded to 65%) was selected as the candidate response threshold for subsequent classification. following PD-1, PD-1+ LAG-3 and PD-1+CTLA4 treatment. **(B)** Cell viability of PDMOs following PD-1, PD-1+ LAG-3 and PD-1+CTLA4 treatment grouped as responder (R), Intermediate, and non-responder (NR). **(C)** Heatmap showing relative organoid viability across all treatment conditions.

**Figure 4.**
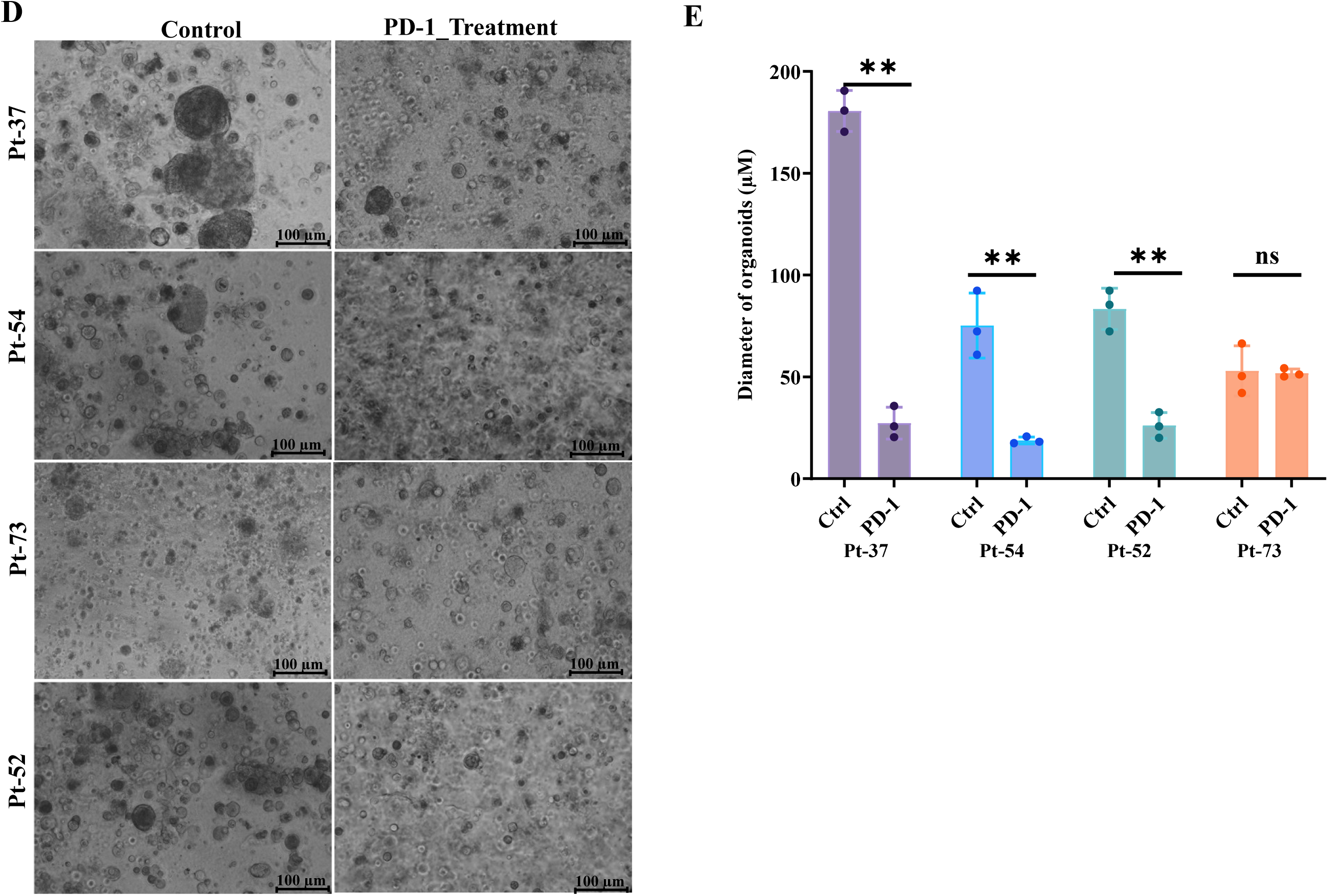
PDMOs morphological response parallels heterogeneous functional sensitivity to PD-1 treatment after day 7. **(D)** Representative bright-field images of PDMOs from four independent patient samples after 7 days of treatment with control and PD-1. Controls display larger and more compact organoids, whereas PD-1 treated organoids show visibly reduced, organoid size and lower structural integrity in all samples. Scale bars: 100 µm. **(E)** Quantification of organoid diameter following 7 day treatment: PD-1 significantly reduced organoid size in P-37 and P-54 (**p < 0.01) but no significant change in P-73 (ns) and induced a moderate but significant reduction in P-52 (\**p < 0.05*). Data represent mean ± SD (n=3).

**Figure 4F.**
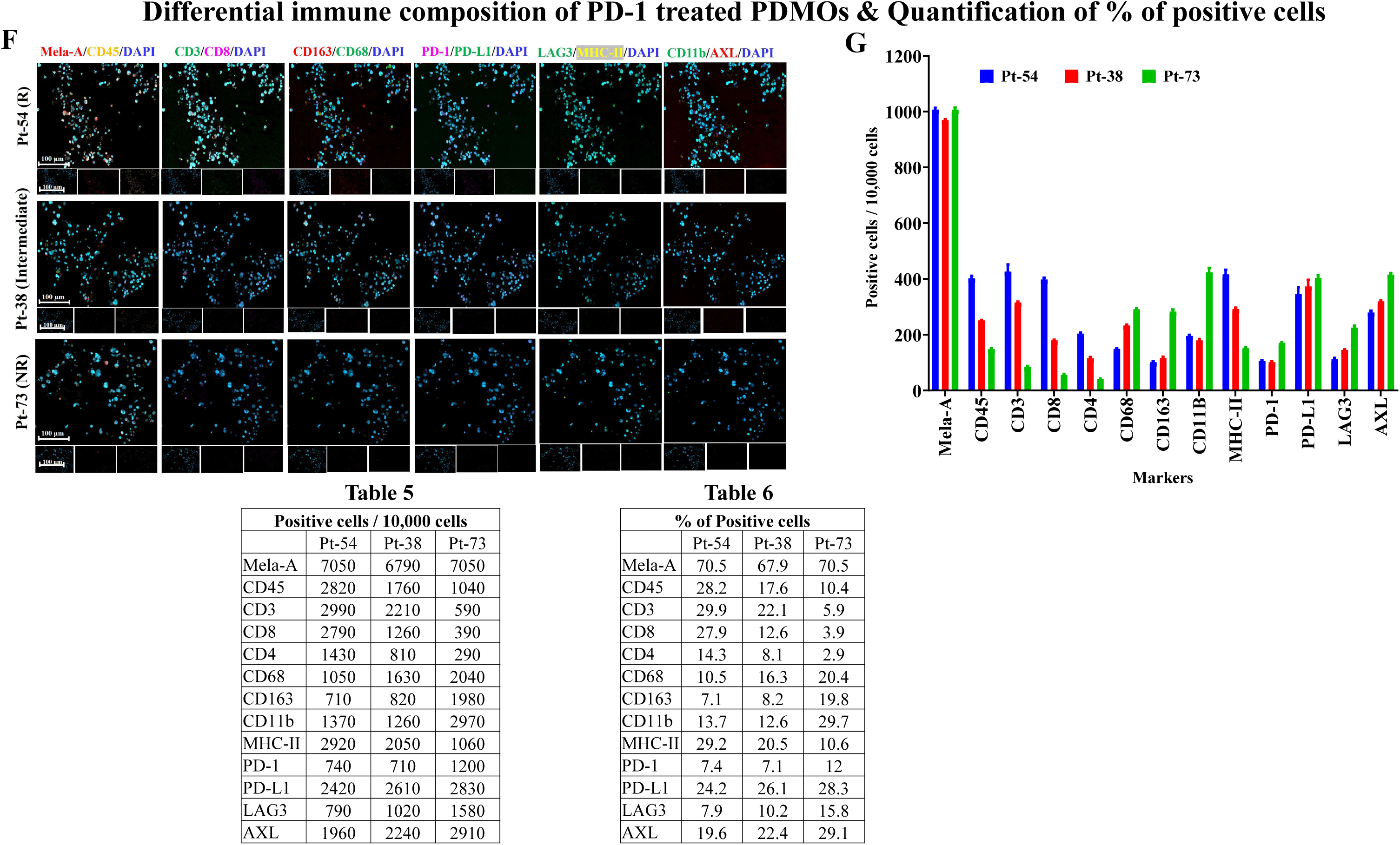
Differential immune composition of PD-1 treated PDMOS after day 7: **(F)** Representative immunofluorescence staining of PDMOs from three patient samples categorized as Responder, Partial Responder and Non-Responder following 7 days of PD-1treatment. Organoids were co-stained for Mela-A (melanoma marker) with immune and phenotypic markers including CD3, CD8, CD11B, CD163, MHC-II, AXL, PD-1, LAG-3, and PD-L1, along with DAPI (nuclei). Responder organoids showed higher infiltration of CD3⁺ and CD8⁺ T cells, increased MHC-II expression, and reduced expression of immunosuppressive markers (AXL). Partial responders demonstrated intermediate levels of T-cell markers and mixed expression of antigen-presentation and suppressive markers. Non-responders showed minimal immune infiltration, low CD8 expression, increased AXL, and persistent PD-1/PD-L1 expression. Insets show individual channels for each marker. Scale bars: 100 µm. Data represent n=3 (3 ROI quantification per organoid). **(G)** Quantification of immune and tumor marker-positive cells per 10,000 cells, demonstrating enhanced T-cell infiltration and antigen presentation in responder organoids and enrichment of immunosuppressive myeloid populations and AXL expression in resistant organoids. Table 5-6: Quantification of % of positive cells on Immunofluorescence in Organoid in difference response with PD-1 mono therapy

**Figure 4H.**
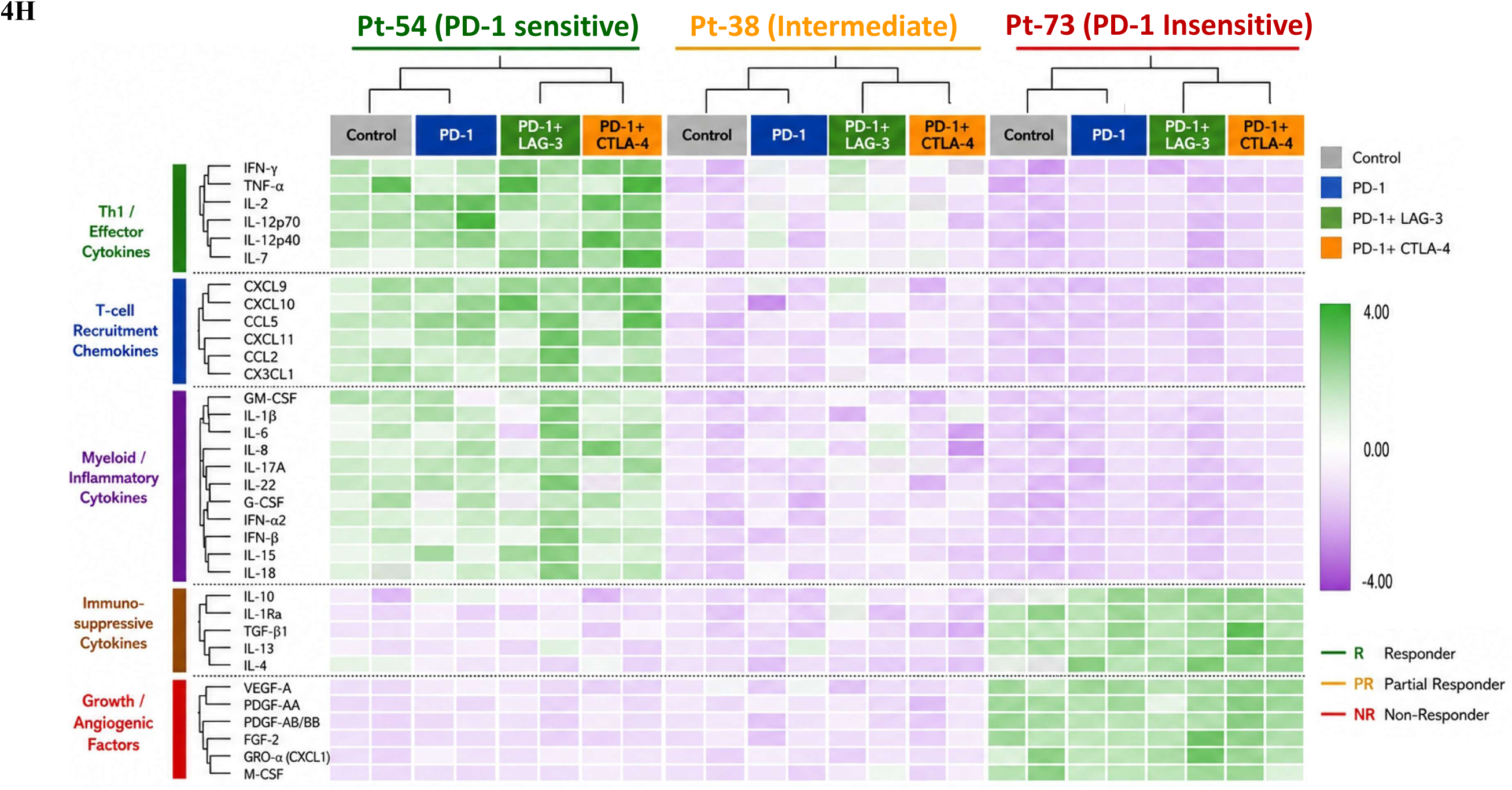
Multiplex cytokine profiling reveals patient-specific immune activation following immune checkpoint blockade. **(I)** hierarchical clustering of multiplex cytokine and chemokine profiles from Pt-54) Pt-38) and Pt-73 patient-derived melanoma organoids following treatment with anti-PD-1, anti-PD-1+anti-LAG-3, or anti-PD-1+anti-CTLA-4. Cytokine expression values are displayed as row-normalized Z-scores, with purple indicating lower expression and green indicating higher expression relative to the mean. PD-1 sensitive organoids exhibited increased expression of Th1-associated inflammatory cytokines and chemokines, including IFN-γ, TNF-α, CXCL9, and CXCL10, particularly following PD-1/LAG-3 blockade. Intermediate responder organoids demonstrated intermediate inflammatory activation, whereas PD-1 insensitive organoids retained elevated expression of immunosuppressive and tumor-associated mediators, including IL-10, IL-1Ra, TGF-β, VEGF-A, PDGF, and FGF-2, with limited induction of effector cytokines. Hierarchical clustering segregated samples according to treatment response, highlighting distinct immune activation states associated with sensitivity or resistance to immune checkpoint inhibition.

**Table 4.** Clinical Treatment and Matched Organoid ICI Responses.

| Clinical vs Organoid ICI responses |  |  |  |
| --- | --- | --- | --- |
| Patients ID's | PD-1 | PD-1+LAG-3 | PD-1+CTLA4 |
| P-52 Clinical | - | - | - |
| P-52 PDMO | R | R | NR |
| P-53 Clinical | - | R | - |
| P-53 PDMO | NR | R | NR |
| P-54 Clinical | - | - | R |
| P-54 PDMO | R | R | NR |
| P-63 Clinical | NR | - | NR |
| P-63 PDMO | R | NR | NR |
| P-74 Clinical | - | R | R |
| P-74 PDMO | R | R | R |
| P-TB7 Clinical | - | - | NR |
| P-TB7 PDMO | R | R | NR |
| P-159 Clinical | - | - | NR |
| P-159 PDMO | Intermediate | Intermediate | Intermediate |
| P-37 Clinical | - | - | - |
| P-37 PDMO | R | R | Intermediate |
| P-77 Clinical | NR | NR | NR |
| P-77 PDMO | NR | NR | NR |
| P-65 Clinical | R | - | - |
| P-65 PDMO | R | R | Intermediate |
| P-10 Clinical | - | NR | - |
| P-10 PDMO | Intermediate | NR | NR |
| P-38 Clinical | - | R | - |
| P-38 PDMO | Intermediate | R | NR |
| P-68 Clinical | NR | - | - |
| P-68 PDMO | NR | Intermediate | R |
| P-164 Clinical | - | - | NR |
| P-164 PDMO | NR | NR | NR |
| P-73 Clinical | - | - | NR |
| P-73 PDMO | NR | NR | NR |

**Note:** R: Responder; NR: Non-Responder

Brightfield imaging following seven days of PD-1 treatment, induced morphological changes and viability responses (Figure 4D, E). Responder organoids, including Pt-37 and Pt-54, exhibited marked reductions in organoid diameter, loss of structural integrity and decreased cellular compaction compared with untreated controls. Pt-52, also a responder, showed smaller absolute reduction in diameter, whereas the non-responder Pt-73 maintained compact morphology with no significant change in diameter following treatment. Quantitative measurements confirmed significant decreases in organoid diameter in responder samples, indicating that morphological alterations represent an additional functional indicator of immunotherapy sensitivity. Post treatment organoids commonly clustered among high-viability groups, supporting their association with acquired resistance following prior anti-PD-1 exposure ^25, 26^.

To investigate the cellular mechanisms underlying these differential responses, immunofluorescence staining was performed on representative responder (Pt-54), intermediate response (Pt-38), and non-responder (Pt-73) organoids following PD-1 treatment (Figure 4F, G, Table 5-6). These features align with prior descriptions of partial clinical responses driven by incomplete T-cell recruitment and tumor cell plasticity^26–28^. Responder organoids displayed abundant infiltration of CD45 and CD3⁺CD8⁺ T cells together with increased MHC-II expression, indicating enhanced immune activation and antigen presentation. In contrast, non-responder organoids exhibited limited lymphocyte infiltration accompanied by enrichment of CD68⁺, CD163⁺, and CD11B⁺ myeloid populations, persistent PD-1/PD-L1 expression, and elevated AXL levels, consistent with an immunosuppressive and therapy-resistant microenvironment. Intermediate responders displayed intermediate immune infiltration together with mixed myeloid and tumor phenotypes. Quantitative analysis (Figure 4G, Table 5-6, Figure S1) confirmed progressive reductions in cytotoxic T-cell infiltration and antigen presentation from responders to non-responders, whereas AXL expression increased with resistance. Collectively, these findings demonstrate that PDMOs faithfully reproduce the functional, morphological and immunological heterogeneity observed in melanoma patients and identify enhanced T-cell infiltration, preserved antigen presentation, and low AXL expression as features associated with responsiveness to immune checkpoint inhibitor. In addition, limited AXL expression suggested a predominance of a melanocytic, therapy-sensitive transcriptional state, as AXL-high mesenchymal programs have been linked to immune evasion and resistance ^28^.

### Multiplex cytokine profiling reveals patient-specific immune activation following immune checkpoint inhibition

To describe the functional immune responses elicited by immune checkpoint inhibition, multiplex cytokine profiling was performed on PD-1 sensitive (Pt-54), Intermediate (Pt-38), and PD-1 Insensitive (Pt-73) patient-derived melanoma organoids following treatment with anti-PD-1 alone and in combination with anti-LAG-3 & anti-CTLA-4 (Figure 4H). Hierarchical clustering demonstrated distinct cytokine expression patterns that separated according to response phenotype, indicating significant inter-patient heterogeneity in immune activation following checkpoint blockade. PD-1 sensitive organoids displayed a matched inflammatory program characterized by increased expression of Th1-associated cytokines, including IFN-γ, TNF-α, IL-2, IL-12p40, IL-12p70, and IL-7, together with enhanced production of T-cell-recruiting chemokines, including CXCL9, CXCL10, CCL5, CXCL11, CCL2 and CX3CL1, consistent with robust activation of adaptive antitumor immunity. Notably, dual PD-1+LAG-3 treatment produced the strongest inflammatory cytokine response, indicating enhanced immune activation compared with PD-1 monotherapy or PD-1+CTLA-4 treatment.

In contrast, Intermediate responder organoids exhibited an intermediate cytokine profile characterized by moderate induction of inflammatory cytokines and chemokines following checkpoint inhibition. Although expression of CXCL9, CXCL10, IL-6, IL-8, IL-12p40/p70, IL-7 and IFN-γ increased relative to untreated controls, the magnitude of induction remained substantially lower than that observed in responder organoids, suggesting incomplete immune reprogramming despite immune checkpoint inhibition. These findings indicate a partially activated tumor immune microenvironment in which immune activation is initiated but remains insufficient to generate a fully effective antitumor response. Such mixed cytokine patterns have been described in partial clinical responders, where immune activation coexists with residual immunosuppressive feedback loops ^29^.

On the other hand, PD-1 Insensitive organoids failed to induce a coordinated inflammatory cytokine following immune checkpoint inhibition and instead maintained an immunosuppressive secretory phenotype. Minimal induction of effector cytokines and T-cell-associated chemokines was accompanied by persistent expression of immunoregulatory mediators, including IL-10, IL-1Ra, TGF-β, IL-4, and IL-13, together with elevated secretion of angiogenic and tumor-promoting factors such as VEGF-A, PDGF-AA, PDGF-AB/BB, FGF-2, CXCL1 and M-CSF, consistent with an immune-excluded, therapy-resistant microenvironment. Across responsive organoids, combination checkpoint blockade elicited stronger inflammatory cytokine responses than PD-1 monotherapy, whereas cytokine profiles in resistant organoids remained largely unchanged irrespective of treatment. Notably, these cytokine signatures closely paralleled the functional treatment responses and immune phenotypes observed in the same organoids (Figures 4A-H), with responder PDMOs exhibiting enhanced CD8⁺ T-cell infiltration, increased MHC-II expression, reduced AXL expression, and robust Th1-associated cytokine production, whereas resistant organoids retained an immune-excluded phenotype characterized by persistent immunosuppressive and angiogenic signaling. In these three models, secreted cytokine profiles paralleled the viability and immunofluorescence phenotypes, suggesting that cytokine profiling may provide a complementary readout whose predictive value remains to be tested.

### Oxygen tension and culture context influence heterogenous responses to immune checkpoint blockade

To examine how culture context and oxygen availability influence immune checkpoint inhibitor (ICI) responses, PDMOs and matched patient-derived two-dimensional (2D) melanoma cultures were treated with anti-PD-1, anti-PD-1+anti-LAG-3, or anti-PD-1+anti-CTLA-4 under normoxic (21% O₂) and reduced-oxygen (5% O₂) conditions. Normalized viability was assessed after seven days of treatment, permitting comparison between culture formats under the same oxygen condition and evaluation of oxygen-dependent changes within individual models (Figure 5; Supplemental Tables 2A and 2B).

**Figure 5.**
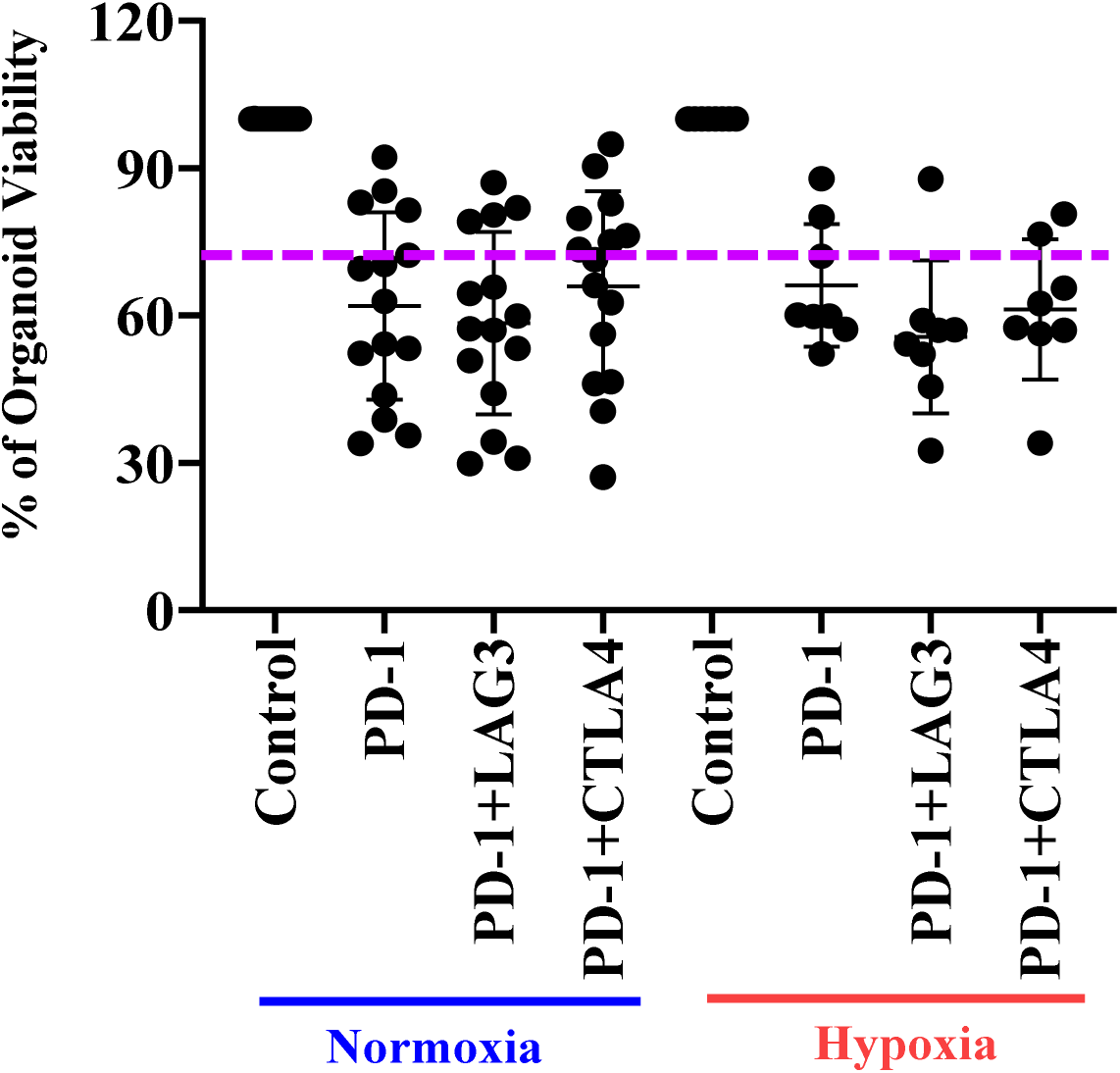
Percent viability of PDMOs following immune checkpoint inhibitor treatment under normoxia (21% O₂) and hypoxia (5% O₂). Values are normalized to untreated controls for each sample. Reduced viability indicates increased sensitivity to ICI treatment. Hypoxia resulted in increased residual viability across the majority of organoids, consistent with hypoxia associated therapeutic resistance. (n = 3).

Under normoxic conditions, PDMOs exhibited heterogeneous checkpoint sensitivities that were not consistently reproduced in matched 2D cultures. Residual viability was frequently lower in PDMOs than in monolayers, although differences between culture formats varied by specimen and treatment, and some model treatment combinations showed similar responses. Thus, cultures derived from the same patient tissue did not necessarily retain equivalent functional sensitivity when maintained in different formats. This inconsistency highlights a fundamental limitation of 2D systems in modeling tumor immune interactions^30,31^.

These differences included clinically responding cases in which treatment sensitivity was more closely reflected by the PDMO. For Pt-74, who responded clinically to PD-1+CTLA-4, residual viability was 46.5% in PDMOs compared with 89.9% in matched 2D cultures. Similarly, Pt-38, who responded clinically to PD-1+LAG-3, showed 57.3% viability in PDMOs compared with 80.9% in monolayers. In these cases, the organoid model retained sensitivity to the clinically active regimen that was attenuated in the corresponding patient-derived monolayer (Table 4; Supplemental Table 3A-3B).

Changing oxygen tension revealed additional variation in checkpoint sensitivity within individual PDMOs. In Pt-37, residual viability following PD-1 blockade increased from 33.9% under normoxia to 80.1% under reduced oxygen, whereas matched 2D cultures showed little change, from 33.3% to 37.1%. The same PDMO also exhibited reduced sensitivity to PD-1+LAG-3 at lower oxygen, with viability increasing from 50.8% to 87.8%; corresponding monolayer values were 50.3% and 45.3%. These comparisons illustrate substantial oxygen-associated losses of checkpoint sensitivity in an individual PDMO that were not reproduced in its matched 2D counterpart (Figure 5, Supplemental Table 3). In this model, reduced oxygen was associated with a substantial loss of checkpoint sensitivity that was not reproduced in the matched monolayer, consistent with reports that hypoxia can limit T-cell-mediated tumor control ^32–33^. Across the 10 PDMOs evaluated under both conditions, residual viable tumor under 5% O_2_ exceeded that under 21% O_2_ in sixteen of thirty model-regimen pairs (PD-1, 7/10; PD-1+LAG-3, 6/10; PD-1+CTLA-4, 3/10), and the direction of change varied by model and regimen.

However, reduced oxygen did not uniformly decrease ICI sensitivity. In Pt-63, PD-1-treated PDMO viability increased from 35.6% to 57.2% at lower oxygen, whereas viability following PD-1+LAG-3 decreased from 80.5% to 52.1%. The corresponding 2D cultures did not reproduce this increased sensitivity to combination blockade, showing viability of 80.8% under normoxia and 87.6% under reduced oxygen. The direction of the oxygen-associated effect therefore differed both between regimens within a PDMO and between matched culture formats.

Together, the baseline differences between culture formats and the oxygen-dependent shifts within individual models demonstrate context-dependent ex vivo checkpoint responses. PDMOs retained clinically relevant sensitivity in selected cases where monolayer responses diverged and revealed environmental modulation that was not consistently reproduced in matched 2D cultures.

## Discussion

Immune checkpoint inhibitors (ICIs) have fundamentally changed the treatment landscape for advanced melanoma; however, considerable interpatient variability and the absence of robust functional biomarkers continue to limit precision immunotherapy. In this study, we established a patient-derived melanoma organoid (PDMO) platform that preserves the cellular complexity of the native tumor microenvironment while enabling rapid ex vivo evaluation of clinically relevant immune checkpoint therapies. Our findings demonstrate that PDMOs retain tumor, stromal, and immune cell populations from the parental tumor, reproduce heterogeneous responses to PD-1-based immunotherapy, and reveal that oxygen tension can modify ex vivo checkpoint sensitivity in a model- and regimen-dependent manner.

A major finding of this study is the high degree of preservation of the native melanoma immune microenvironment. Unlike conventional two-dimensional cultures, which rapidly lose stromal and immune components, PDMOs maintained melanoma lineage markers together with lymphoid, myeloid, stromal, and immune checkpoint populations. The preservation of CD3⁺, CD8⁺, CD68⁺, CD163⁺, CD11B⁺, PD-1, and PD-L1 expression indicates that the organoid system retains many of the cellular interactions required to model immune checkpoint responses. The close agreement between matched parental tumors and organoids for the principal populations, with quantitative differences for a subset of markers, extends previous observations that melanoma organoids maintain molecular and histopathological features of their originating tumors. The maintenance of these populations may contribute to the divergent checkpoint sensitivities observed between PDMOs and matched monolayers, although predictive capacity was not formally compared.

Another important observation is the association between NGFR (CD271) expression and successful organoid establishment. Tumors enriched for NGFR-positive melanoma cells consistently generated stable long-term organoid cultures, whereas specimens with low NGFR expression frequently exhibited limited growth or failed to establish altogether. NGFR has previously been implicated in melanoma stemness, cellular plasticity, tumor initiation, and therapeutic resistance. NGFR-positive melanoma cells have been previously linked to a neural crest like, dedifferentiated state characterized by stress tolerance, plasticity, and therapy resistance ^10^. Our findings suggest that NGFR may additionally identify melanoma cell populations capable of sustaining three-dimensional growth ex vivo. Although these observations require validation in larger patient cohorts, they provide a practical biomarker for predicting organoid establishment and further support the biological importance of NGFR-positive melanoma subpopulations.

Functional immune checkpoint inhibitor screening demonstrated substantial interpatient heterogeneity, emphasizing the need for individualized therapeutic assessment. Anti-PD-1 monotherapy distinguished responder, intermediate, and non-responder organoids. Neither combination regimen produced a uniform reduction in viability relative to PD-1 monotherapy; the regimen associated with the lowest residual viability differed among models, consistent with patient-specific rather than regimen-wide sensitivity. This pattern is compatible with the clinical experience that the incremental benefit of adding anti-LAG-3 or anti-CTLA-4 to PD-1 blockade, as established in RELATIVITY-047 and CheckMate 067 ^2,3^, is realized in a subset of patients. Organoid calls agreed with available clinical outcomes in fourteen of seventeen patient-regimen pairs, with two discordant cases, supporting further evaluation of functional ex vivo testing as a complement to existing molecular biomarkers, with the caveat that the candidate threshold was derived from the same outcomes. Rather than relying solely on static biomarker measurements such as PD-L1 expression or tumor mutational burden, PDMOs integrate dynamic tumor-immune interactions into a functional readout of therapeutic sensitivity. Prior studies have demonstrated that the presence, spatial distribution, and functional state of tumor-infiltrating lymphocytes are critical determinants of response to ICIs in melanoma^34–35^.

Immune profiling of three models showed that therapeutic responsiveness was associated with coordinated differences in immune composition and secreted cytokines. Responsive organoids exhibited increased CD3⁺ and CD8⁺ T-cell infiltration, enhanced MHC-II expression, reduced AXL expression, and robust induction of inflammatory cytokines and chemokines, including IFN-γ, TNF-α, IL-2, CXCL9, CXCL10, and CCL5. In contrast, resistant organoids maintained persistent PD-L1 expression together with enrichment of CD68⁺/CD163⁺ myeloid populations and elevated expression of AXL, accompanied by immunosuppressive and angiogenic cytokines. These findings are consistent with current models of melanoma immune resistance, in which dedifferentiated AXL-high melanoma cells, suppressive myeloid populations, impaired antigen presentation, and deficient T-cell recruitment collectively contribute to failure of immune checkpoint blockade. This intra-tumoral plasticity has been recognized as a major driver of resistance to both targeted and immunotherapies^36^. The close agreement between cellular phenotypes and cytokine signatures suggests that multiplex cytokine profiling may serve as a complementary functional biomarker capable of monitoring treatment-induced immune activation within patient-derived organoids.

The oxygen comparisons highlight the importance of experimental context when interpreting ex vivo checkpoint responses. Lower oxygen was not associated with uniform resistance; instead, the direction and magnitude of the observed viability changes varied across patient-derived models and therapeutic regimens. Differences between PDMOs and matched two-dimensional cultures further suggest that treatment responses depend on the culture system, although this comparison does not isolate tissue architecture from other differences that may arise between the models.

These experiments do not establish whether oxygen-associated viability changes reflect altered tumor-cell behavior, immune-cell activity, cellular composition, or other contributions to the assay readout. Rather, they identify a setting in which those contributions can be investigated. The findings also motivate the hypothesis that local environmental differences may contribute to heterogeneous treatment responses among lesions within an individual patient. That clinical mechanism was not directly tested here, but the ability to examine the same patient-derived model under different environmental conditions provides a tractable approach for future investigation. Whether the oxygen-associated shifts observed ex vivo relate to intratumoral hypoxia and clinical outcome, as proposed for hypoxic melanomas ^37–38^, was not examined here.

This study has several limitations. First, although the organoid establishment rate was encouraging, successful long-term cultures were obtained from only 60% of collected specimens, indicating that optimization of culture conditions remains necessary for broader clinical implementation. Second, the number of organoids subjected to detailed functional immune profiling was relatively limited, and larger prospective cohorts will be required to validate predictive biomarkers identified in this study. Third, although PDMOs preserve endogenous immune populations during early culture, immune cell viability declines with prolonged maintenance, limiting long-term investigations of adaptive immune responses. Incorporation of autologous peripheral blood mononuclear cells, tumor-infiltrating lymphocytes, or additional stromal components may further improve physiological relevance. Finally, correlation with larger prospective clinical response datasets will be essential before implementation as a clinical companion diagnostic platform.

Despite these limitations, our study provides initial evidence that patient-derived melanoma organoids represent a biologically relevant and translationally valuable model for precision immuno-oncology. The ability to preserve patient-specific tumor heterogeneity while enabling rapid functional testing of immune checkpoint inhibitors offers significant opportunities for individualized treatment selection, biomarker discovery, and mechanistic investigation of immunotherapy resistance. Future integration of organoid-based functional profiling with genomic, transcriptomic, and spatial immune analyses may further enhance prediction of therapeutic response and facilitate rational development of personalized combination immunotherapies.

## Conclusions

We established an early-passage PDMO platform that retains key tumor, immune, and stromal components of parental melanoma and permits functional examination of patient-derived tumor biology. PDMOs exhibited heterogeneous responses to checkpoint blockade across patient-derived models, therapeutic regimens, and oxygen conditions. Selected models also displayed distinct immune-cell and cytokine profiles, providing complementary cellular and secretory characterization. These findings support PDMOs as an ex vivo model for investigating tumor-immune interactions and context-dependent treatment sensitivity. The platform provides a basis for further studies of how endogenous cellular composition and environmental conditions contribute to heterogeneous immunotherapy responses.

## Acknowledgements

The authors thank the Stanford Human Immune Monitoring Center for assistance with Luminex cytokine profiling and the patients who generously contributed tumor samples to this study.

## Authors’ Contributions

MS conceived the study, performed experiments, organoid generation, analyzed data and wrote the manuscript. Co-authors assisted with organoid generation, immunofluorescence analysis, cytokine profiling and data interpretation. AK supervised the study and critically revised the manuscript. All authors read and approved of the final manuscript.

## Funding

This work was generously supported by the John and Marva Warnock Endowed Scholar Fund and the Melanoma Research Alliance Team Science Award (Award #)

## Competing Interests

The authors declare that they have no competing interests.

## Patient consent

Written informed consent was obtained from all participants prior to sample collection.

## Ethics Approval

Human melanoma specimens were obtained under an Institutional Review Board and approved protocol at Stanford University (IRB #65607).

## Availability of Data and Materials

All data relevant to the study are included in the article and its supplementary information files. Additional datasets are available on reasonable request.

## Supplemental Figures and Tables

**Table 1-3:** Quantification of % of positive cells on Immunofluorescence in Organoid vs matched Parental tissue and their p values.

**Table 4.** Clinical Treatment and Matched Organoid ICI Responses

**Table 5-6:** Quantification of % of positive cells on Immunofluorescence in Organoid in difference response with PD-1 mono therapy

**Supplemental Table 1.** Clinical parameters of selecting patients’ information

**Supplemental Table 2A:** Percent viability of 3D following immune checkpoint inhibitor treatment under hypoxic conditions: Values are normalized to untreated controls.

**Supplemental Table 2B:** Shows viability changes of matched patient-derived melanoma cells cultured in 2D following immune checkpoint inhibitor treatment under normoxia and hypoxic conditions: Values are normalized to untreated controls. (n = 3)

**Supplemental Table 3 A:** Shows viability changes of PDMOs following immune checkpoint inhibitor treatment under normoxia (21% O₂) and hypoxia (5% O₂). Values are normalized to untreated controls for each sample. Reduced viability indicates increased sensitivity to ICI treatment. Hypoxia resulted in increased residual viability across the majority of organoids, consistent with hypoxia associated therapeutic resistance. (n = 3) *Note: “-” Indicates not growing in hypoxic conditions*.

**Supplemental Table 3B:** Shows viability changes of matched patient-derived melanoma cells cultured in 2D following immune checkpoint inhibitor treatment under normoxia and hypoxic conditions: Values are normalized to untreated controls. Compared with 3D organoids, 2D cultures confirmed that consistently higher viability and minimal oxygen-dependent modulation, indicating limited sensitivity to immune checkpoint inhibition and reduced modeling of hypoxia-induced resistance (n = 3). Note: Red: indicates significant difference between 2D vs 3D normoxia; Blue: indicates significant difference between 2D vs 3D Hypoxia: Brown: indicates significant difference in 3D normoxia vs Hypoxia

**Supplemental Table 4A-C:** Quantification of % of positive cells on Immunofluorescence in Organoid vs matched Parental tissue and their p values.

**Table S1.** Clinical parameters of selecting patients information.

| Clinical Information |  |  |  |  |  |
| --- | --- | --- | --- | --- | --- |
| Therapy | Patients | Age | Sex | Stag | Mutational status |
| Primary | Pt-52 | 72 | F | IIC |  |
| Pre | Pt-53 | 57 | F | III | BRAF, NRAS,<br>NED |
| Primary | Pt-54 | 77 | M | IIC | NED (CR), BRAF |
| TIL | Pt-63 | 63 | F | IV | BRAF WT,<br>NARAS |
| TIL | Pt-74 | 64 | F | IV | BRAF WT |
| Primary | Pt-TB7 | 60 | F | IV |  |
| TIL | Pt-159 | 41 | M | IV | BRAF (+) |
| Primary | Pt-37 | 60 | F | IIA |  |
| Primary | Pt-77 | 70 | F | IV | BRAF WT |
| Pre | Pt-65 | 71 | M | IIIB | BRAF (+) |
| Post | Pt-10 | 73 | M | IIC | BRAF WT |
| Post | Pt-38 | 71 | M | III D | BRAF WT |
| Post | Pt-68 | 60 | F | IIIB | BRAF (+) |
| Post | Pt-164 | 52 | M | 1A | BRAF (+) |
| TIL | Pt-73 | 54 | M | IV | BRAF (+) |
| Primary | Pt-112 | 83 | M | IIC |  |
| Pre | Pt-166 | 65 | M | IIIC |  |

**S Table 2A)** Percent viability of 3D following immune checkpoint inhibitor treatment under hypoxic conditions: Values are normalized to untreated controls. **S Table 2B)** Percent viability of matched patient-derived melanoma cells cultured in 2D following immune checkpoint inhibitor treatment under normoxia and hypoxic conditions: Values are normalized to untreated controls. (n = 3)

**Supplemental Table 2A**
| Patients ID's | **3D_ viability loss/improved response under hypoxia** |  |  |
| --- | --- | --- | --- |
|  | **PD-1** | **PD-1+LAG3** | **PD-1+CTLA4** |
| Pt-52 | 16.17 | 0.97 | -17.39 |
| Pt-53 | -4.36 | 22.65 | -13.93 |
| Pt-54 | 13.42 | -2.81 | -7.83 |
| Pt-63 | 21.59 | -28.31 | 5.71 |
| Pt-74 | 6.82 | 2.73 | 9.77 |
| Pt-TB7 | 5.53 | 14.64 | 10.95 |
| Pt-159 | 2.51 | -5.48 | -22.18 |
| Pt-37 | 46.22 | 37.02 | 14.05 |
| Pt-77 | NA | NA | NA |
| Pt-65 | NA | NA | NA |
| Pt-10 | NA | NA | NA |
| Pt-38 | NA | NA | NA |
| Pt-68 | NA | NA | NA |
| Pt-164 | NA | NA | NA |
| Pt-73 | NA | NA | NA |

**Supplemental Table 2B**
| Patients ID's | **2D_ viability loss/improved response under hypoxia** |  |  |
| --- | --- | --- | --- |
|  | **PD-1** | **PD-1+LAG3** | **PD-1+CTLA4** |
| Pt-52 | 5.19 | -1.96 | -9.14 |
| Pt-53 | 2.10 | 6.48 | 3.59 |
| Pt-54 | 2.67 | -2.14 | -2.54 |
| Pt-63 | 1.62 | 6.76 | -3.80 |
| Pt-74 | -2.55 | 6.02 | -5.18 |
| Pt-TB7 | 11.22 | 9.23 | -5.82 |
| Pt-159 | 14.53 | 10.29 | -18.58 |
| Pt-37 | 3.81 | -5.01 | 11.33 |
| Pt-77 | -2.18 | -0.42 | -2.76 |
| Pt-65 | -6.76 | 21.32 | 6.26 |
| Pt-10 | 20.01 | 5.21 | 0.21 |
| Pt-38 | 10.72 | 2.92 | -7.29 |
| Pt-68 | 4.65 | -15.03 | 8.74 |
| Pt-164 | 19.76 | 1.01 | 1.91 |
| Pt-73 | 1.54 | -4.09 | -5.26 |

**Supplemental Table 3.**
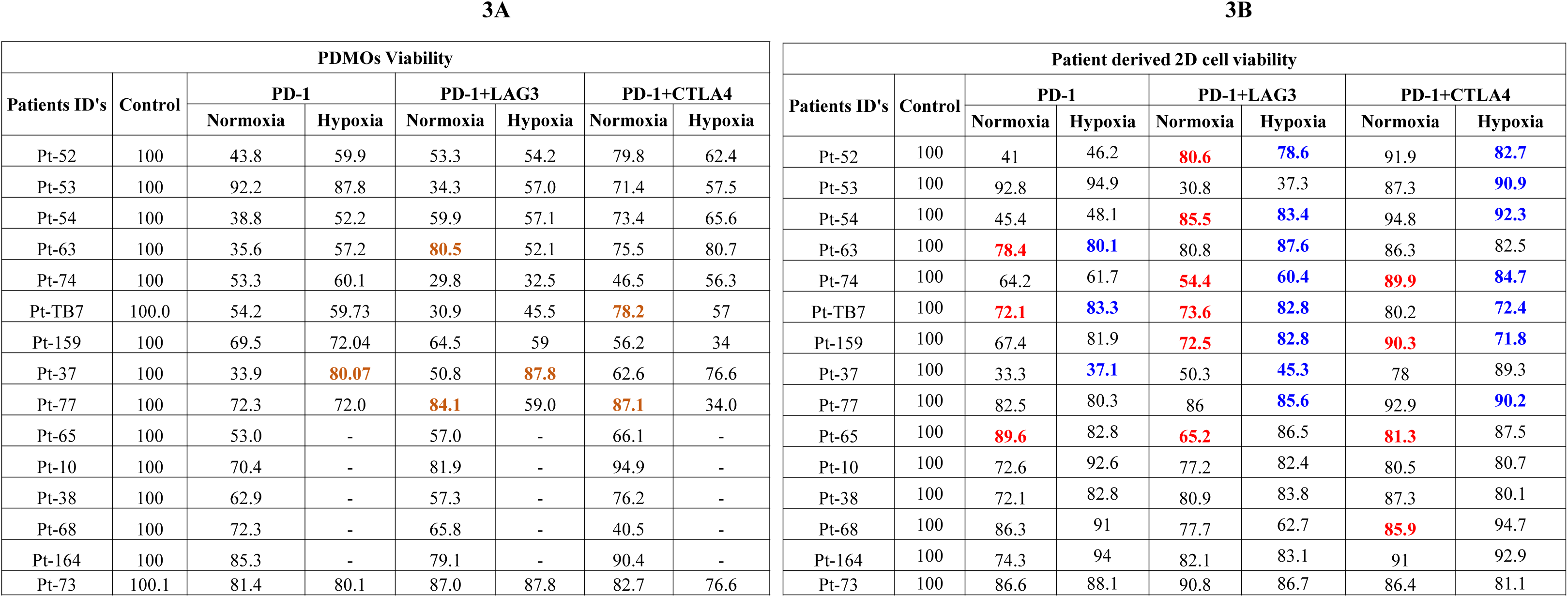
A) Percent viability of PDMOs following immune checkpoint inhibitor treatment under normoxia (21% O₂) and hypoxia (5% O₂). Values are normalized to untreated controls for each sample. Reduced viability indicates increased sensitivity to ICI treatment. Hypoxia resulted in increased residual viability across the majority of organoids, consistent with hypoxia associated therapeutic resistance. (n = 3) *Note: “-” Indicates not growing in hypoxic conditions*. **B)** Percent viability of matched patient-derived melanoma cells cultured in 2D following immune checkpoint inhibitor treatment under normoxia and hypoxic conditions: Values are normalized to untreated controls. Compared with 3D organoids, 2D cultures confirmed that consistently higher viability and minimal oxygen-dependent modulation, indicating limited sensitivity to immune checkpoint inhibition and reduced modeling of hypoxia-induced resistance (n = 3). Note: **Red**: indicates significant difference between 2D vs 3D normoxia; **Blue**: indicates significant difference between 2D vs 3D Hypoxia: **Brown:** indicates significant difference in 3D normoxia vs Hypoxia

**Supplemental Figure 1:**
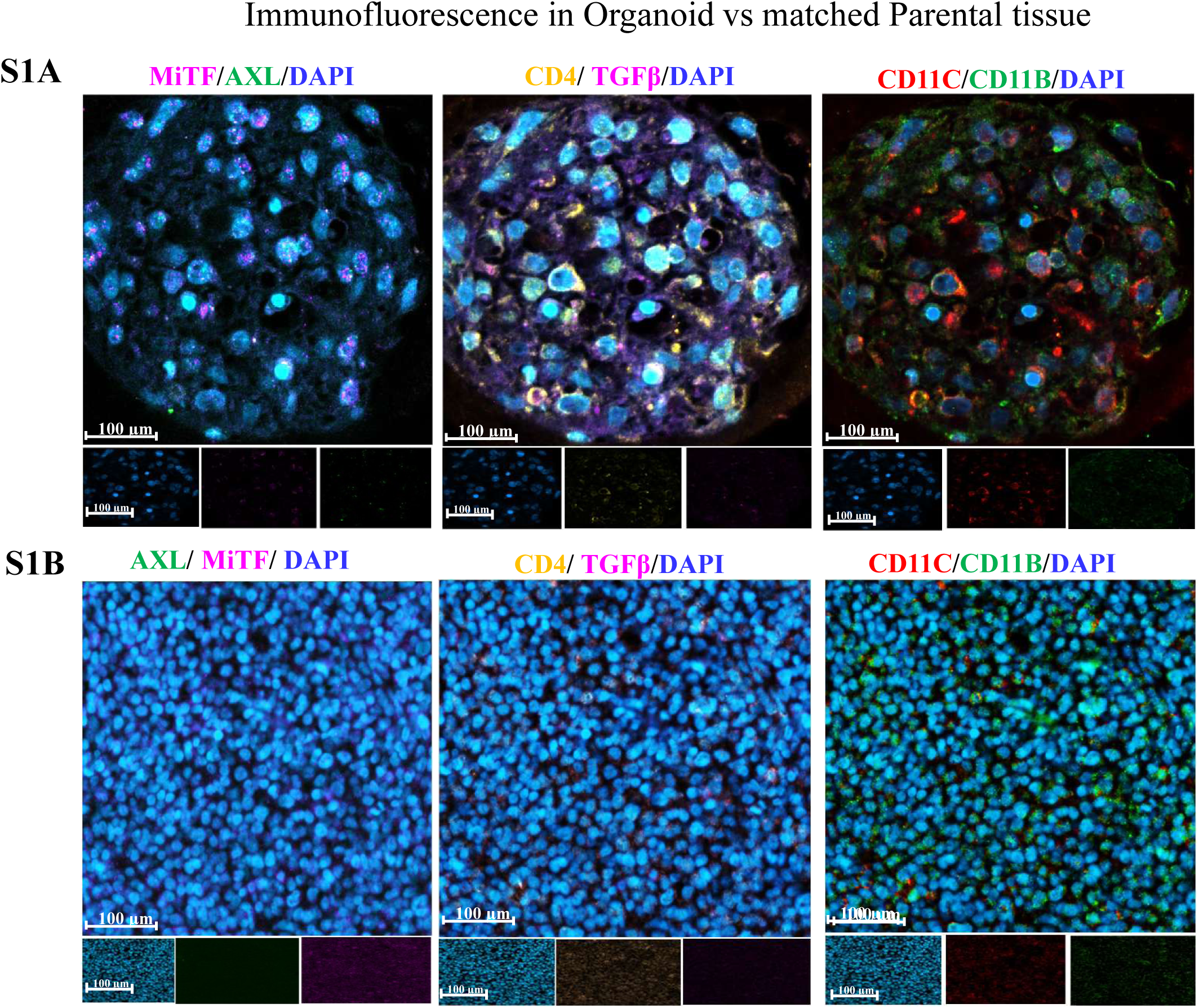
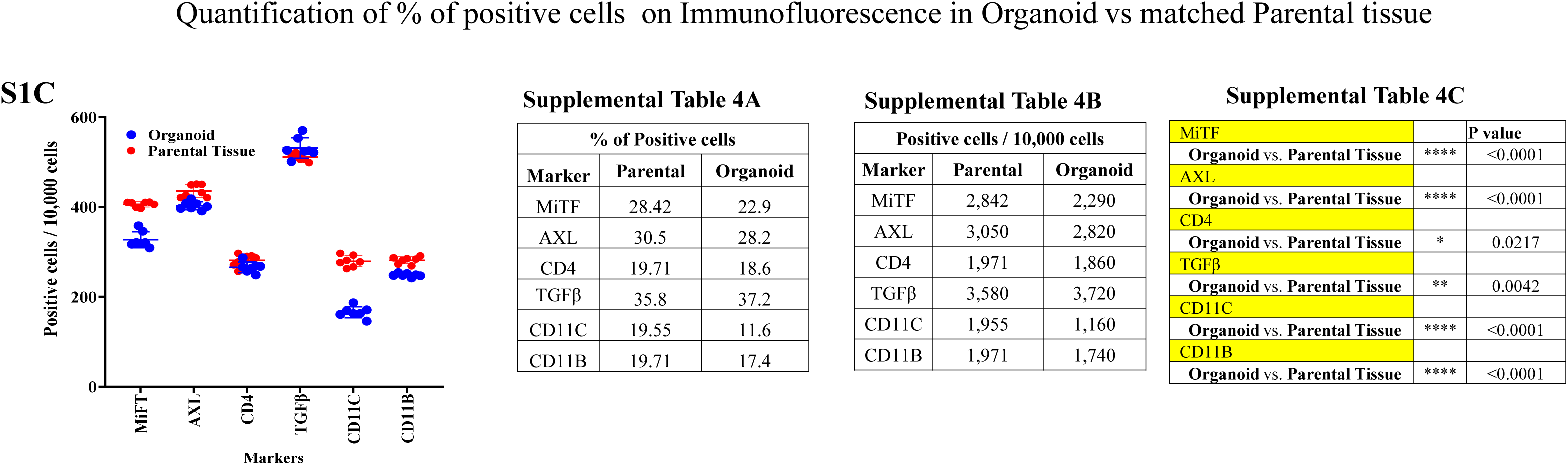
Immunofluorescence characterization of PDMOs and matched parental tumor tissue. **(S1A)** PDMOs immunostaining: Representative immunofluorescence images of melanoma organoids stained for immune and tumor-associated markers: CD4, CD11B, CD11C, MHC-II, MITF, AXL and TGFβ, Nuclei are stained with DAPI, Insets show single-channel images for each marker. Scale bar: 100 µm. These data demonstrate the heterogeneous composition of the PDMOs and the presence of key immune and tumor cell populations within the 3D system. **(S1B)** Matched parental tissue Immunofluorescence stained with the same antibody panel. Scale bars: 20 µm (overview), 100 µm (zoomed panels). Marker expression patterns in the parental tumor mirror those observed in the organoids. Comparative analysis of panels D and E demonstrates that the PDMOs faithfully recapitulate the cellular heterogeneity, immune and expression of melanoma differentiation, dedifferentiation and immune checkpoint markers found in the original parental tissue. **(S1C)** Quantitative comparison of marker-positive cells in matched parental tumors and PDMOs. Comparable expression profiles were observed across immune and immune checkpoint. Data are presented as mean ± SD. Data represent n=7 (3 ROI quantification per organoid).

## Notes

### Competing Interest Statement

The authors have declared no competing interest.

